# An Integrative Approach to Rational Engineering of Dengue Virus-Like Particles

**DOI:** 10.1101/2025.03.11.642607

**Authors:** Venkata Raghuvamsi Palur, Fan-Chi Chen, Shang-Rung Wu, Guan-Wen Chen, Peter J. Bond, Day-Yu Chao, Jan K. Marzinek

## Abstract

Virus-like particles (VLPs) are promising vaccine candidates due to their noninfectious and highly immunogenic nature. These particles lack a viral genomic core and display a robust host immune response. VLPs are typically highly unstable and heterogeneous in size. This motivates the characterization of the biophysical and structural properties of VLPs to facilitate the rational design of stable and highly immunogenic particles. We employed an integrative approach combining multiscale modeling, lipidomics, and *in vitro* experiments to gain molecular insights into the factors governing VLP stability and homogeneity. We focused on dengue virus VLPs, which are known to elicit neutralizing antibodies similar to infectious virions. Systematic introduction of mutations in the structurally crucial stem helix region of the chimeric E protein guided by molecular simulations allowed us to modulate the secretion efficiency of VLPs *in vitro*. Overall, this work highlights the role of protein□lipid envelope interactions in maintaining VLP stability with better yield, providing a framework for the future development of stable and immunogenic next-generation VLP-based vaccines.

## Introduction

DENV is an enveloped virus that infects millions of people worldwide and significantly impacts human life and economies[1, 2]. DENV, which belongs to the *Flaviviridae* family and was recently renamed *Orthoflavivirus*[3], includes other serious human pathogens, such as Zika virus (ZIKV), Japanese encephalitis virus (JEV), West Nile virus, and yellow fever virus (YFV). The outer layer of the ∼50 nm diameter virion is made of 180 copies of glycosylated envelope (E) and membrane (M) proteins embedded in the lipid bilayer, which encapsulate the ∼10 kbp RNA genome in complex with the capsid protein. The ectodomain of the E protein contains three domains (DI, DII, and DIII), which lie flat on the viral membrane in the mature “smooth” virus. This sequence is followed by an α-helical stem region composed of three stem helices (E-H1, E-H2, and E-H3) and a C-terminal transmembrane (TM) region containing two TM helices (E-T1 and E-T2). Immature DENV consists of 60 trimeric E and precursor membrane (prM) proteins, which form “spikes” emerging from the viral surface. In this state, each E protein hydrophobic fusion peptide is capped with a pr molecule, which prevents premature fusion of the virus. During the maturation process in the cell at low pH, accompanied by host furin protease-mediated prM cleavage, large-scale conformational changes in E proteins lead to a switch from noninfectious immature trimeric spikes to infectious mature dimers in smooth mature virions. Finally, mature DENV particles are secreted into the extracellular space.

There are four DENV serotypes whose individual surface proteins and virus assembly are nearly identical, and they share a sequence similarity of ∼65-70% at the entire genome level. During the viral life cycle, due to environmental factors such as pH, temperature, or the presence of divalent cations, the virus adopts numerous conformations with varying exposure of epitope sites[4, 5]. Collectively, these findings complicate efforts to generate antibodies that neutralize all serotypes and viral morphologies[6] and can also result in antibody-dependent enhancement (ADE), making it a challenge to develop highly potent and safe vaccines[7–9]. Unfortunately, current vaccines have not accounted for both challenges. For instance, Dengvaxia®, developed by Sanofi Pasteur, has low efficacy and is associated with dangerous health outcomes among younger people or people without prior dengue infections after vaccination[10, 11]. More recently, a tetravalent live attenuated vaccine developed by Takeda Pharmaceuticals (QDENGA, TAK-003) was approved in Brazil for people regardless of prior dengue exposure[12, 13]. Still, it was reported to show varying and waning efficacy among serotypes[14]. These factors make dengue vaccine development a major challenge from both an immunological and a structural point of view. Thus, there is an urgent need for novel and effective therapeutics.

VLPs represent promising platforms for vaccine alternatives because they lack genomic material and hence are noninfectious, a key factor in vaccine safety[15]. VLPs are nanoparticles whose structures resemble those of viruses, and can trigger significant immune responses as shown by multiple FDA-approved vaccines and candidates in clinical trials [16]. Like viruses, VLPs can be classified as nonenveloped VLPs (neVLPs) or enveloped VLPs (eVLPs)[17]. Unlike eVLPs, engineering neVLPs has achieved considerable success by manipulating the selectivity of cargo, stability, and homogeneity in particle assemblies[18–20]. The mature DENV serotype 2 (DENV-2) VLP (mD2VLP) was previously reported to induce broad neutralizing antibodies against all four DENV serotypes. A more recent study of a tetravalent dengue VLP-based vaccine revealed that the vaccine generates neutralizing antibodies against all serotypes in nonhuman primates[23]. In contrast to dengue virions, mature dengue VLPs are only ∼26-36 nm in diameter and are composed of 30 E and M protein dimers embedded within lipid vesicles. Previously, the cryo-electron microscopy (cryoEM) density map of mD2VLP was solved at ∼13 Å resolution. The mD2VLP particle was shown to be a T=1 icosahedral with a smooth surface. However, due to the absence of RNA and capsid proteins, VLPs are highly heterogeneous in morphology. With the current low resolution of the cryoEM mD2VLP data and missing structural details such as secondary structure at the protein-lipid envelope interface as well as the location of the stem helix/TM regions in lipid envelope, these pose a challenge to engineering DENV VLPs for desired physiochemical properties with optimal protein-lipid interactions.

Extensive efforts have been directed toward engineering VLPs with desirable biophysical and immunological properties. For instance, chimeric VLPs were previously generated with sequences corresponding to the ectodomain of DENV-2 and stem-TM regions of either JEV or vesicular stomatitis virus to increase the secretion and production of VLPs[24, 25]. High secretion of dengue and Zika VLPs in mammalian cells was previously shown via an F108A mutation and by replacing the stem/TM region of the mutation corresponding to DENV-1[26]. Although VLPs possess significant advantages over conventional vaccine strategies, their low secretion and low stability are serious drawbacks that require sequence modifications.

In this work, we integrate experimental and computational techniques to identify critical protein lipid interactions to engineer VLPs with improved secretory properties. We focused on mD2VLP, which has been shown to lead to highly immunogenic responses in mice. First, we performed mass spectrometry-based lipidome profiling to determine the lipid envelope composition of VLPs from different cell lines. Guided by the cryo-EM density map of mD2VLP, we constructed all-atom (AA) models of the EM complex, and performed extensive molecular dynamics (MD) simulations to probe the protein lipid and protein protein interactions between E and M dimers governing particle stability..*In silico* mutations that predicted stronger protein lipid interactions were validated by site-directed mutagenesis experiments in which substantial increases in VLP secretion were observed. Finally, coarse-grained (CG) simulations of the dengue VLPs reproduced particle morphologies that closely resembled cryo-EM micrographs, while the observed dynamics revealed the basis for VLP heterogeneity in solution. Overall, this work lays a general foundation for the rational development of highly secreting and effective VLPs for the use in next-generation vaccines.

## Results

### The lipid composition of VLPs is conserved across flaviviruses and cell lines

We first sought to characterize the lipid content of VLPs derived from several flaviviruses under different conditions. We produced mD2VLPs from different cell lines, and the lipidomic profiles of VLPs from each cell line were identified. Three different purified VLPs were used: i) dengue mD2VLP prepared from both HEK-293T and COS-1 cells; ii) JEV VLP (JEVLP) prepared from COS-1 cells; and iii) ZIKV VLP (ZIKAVLP) prepared from COS-1 cells. The lipids were subjected to lipid extraction followed by liquid chromatographic tandem mass spectrometry (LC MS/MS) to obtain lipidomic data after confirmation of particle formation by electron microscopy (EM) analysis (Fig. 1A-D). Detailed descriptions of VLP production, purification, lipid extraction, and LC MS/MS can be found in the Methods section. The results of LC MS/MS analysis revealed similar lipid compositions across different flavivirus VLPs from the same cell lines or the same VLPs from different cell lines. The two types of lipids extracted in the highest quantity from purified VLPs were diacylglycerols (DGs) in the glycerolipid category, and fatty acids (FAs) in the fatty acyls category; these are mainly unsaturated fatty acid chains with DG oxidized on carbon number 34 and FAs oxidized on carbon number 18. Phosphatidylcholine (PC), a glycerophospholipid; oxidized fatty acids (OxFAs), a fatty acyl; and other lipids, such as ceramide alpha-hydroxy fatty acid-phytospingosine (cerAP), in the sphingolipid category, had relatively low abundances (Fig. 1E). By summing all the *m/z* spectra of the positive or negative lipids, no differences in the total amount of lipids were observed among the different types of VLPs (Fig. 1F-G).

**Fig. 1.**
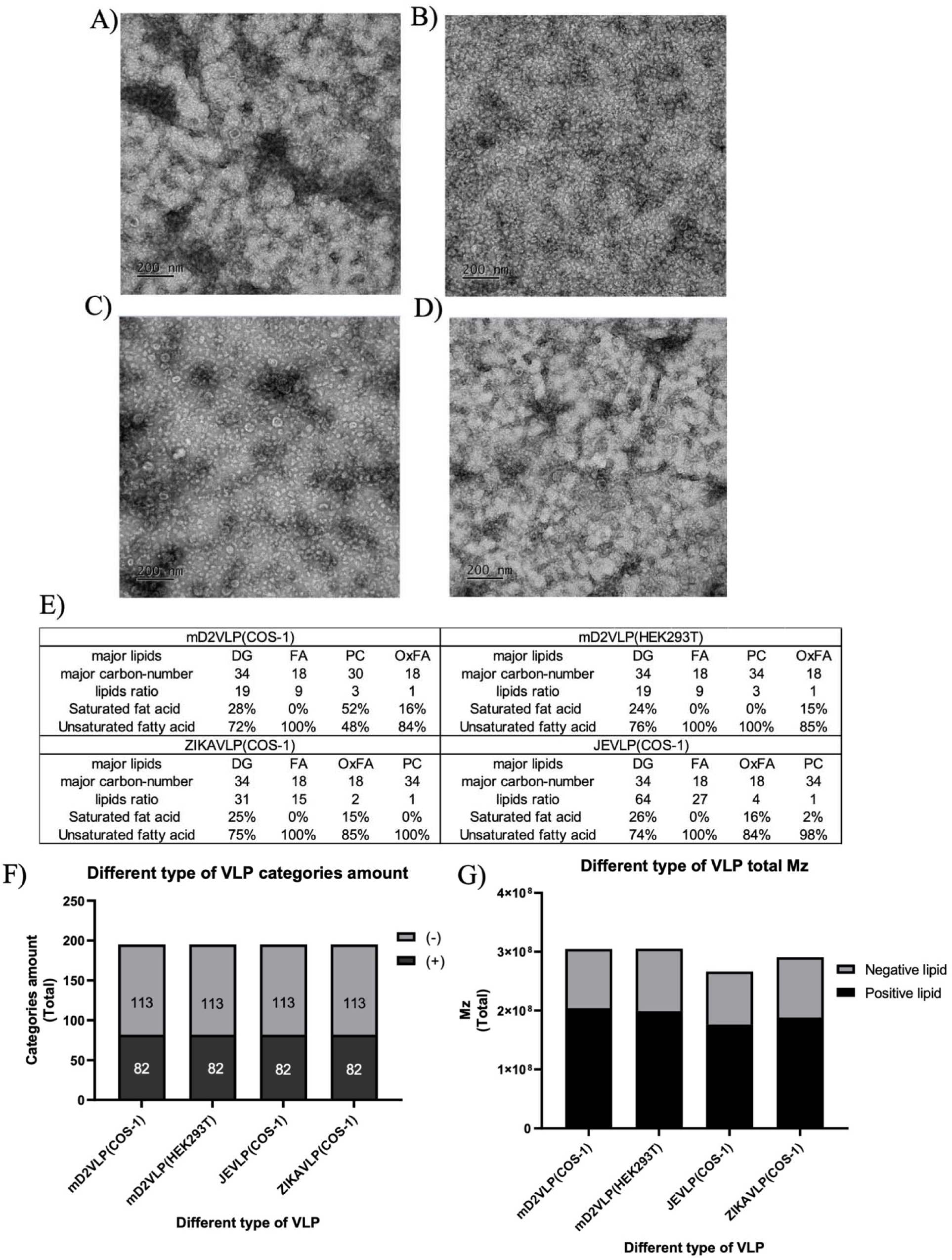
Lipidomic analysis of four different flavivirus virus-like particles (VLPs) via liquid chromatography (LC) tandem mass spectrometry analysis. **(A-D)**. Representative electron microscopy images examined after flavivirus VLP purification from **(A)** mature DENV-2 VLPs (mD2VLPs) secreted from stable HEK293T cells and **(B)** Japanese encephalitis virus VLPs (JEVLP), (C) mD2VLPs and (D) Zika virus VLPs (ZIKAVLPs) generated from transiently transfected COS-1 cells. **(E)** The four highest quantities of lipids extracted from different VLPs. **(F)** The total amount of all lipid categories detected from different purified VLPs based on positive (+) and negative (-) ion modes. **(G)** The total M/z ratio of positive or negative lipids of all lipid categories detected from different purified VLPs. No differences in total amount or M/z were found among the different VLPs. DG: diacylglycerols; FA: fatty acid; PC: phosphatidylcholine; OxFA: oxidized fatty acid.

### Specific E protein**□**lipid interactions govern VLP secretion efficiency

Previous studies have shown that the secretion of mature DENV-2 VLPs following the transfection of prM/E plasmids into mammalian cells can be achieved only with an E protein chimera (Echi), in which residues 395-495 of DENV-2, which represents part of the stem helix-transmembrane (TM) region, are replaced with JEV[27] (Fig. 2A). Through helical wheel analysis of E-H1, we identified several amino acid residues that we hypothesized may play a crucial role in VLP stability and secretion via interactions with the lipid envelope. In particular, dimeric E-M protein structures revealed that residues L398, F402, T405, A409, L412, and L415 all faced the lipid bilayer (Fig. 2B). To assess the contribution of these residues to VLP secretion, we performed site-directed mutagenesis at each site to alanine (A) or glutamate (E) to interrupt the interaction with the lipid bilayer (E-H1 mutants). The rationale of choosing A or E were to probe the effect of side chain removal or introducing a negative charge at the negatively charged lipid head group region, both of which can potentially lower VLP secretion. The mutant plasmids were transfected into HEK293T cells to investigate changes in the relative VLP secretion efficiency with respect to that of the wild type through antigen-capture ELISA. The results revealed that all the mutations had negative effects on VLP secretions. Among them, mutations at A409E and L412A/E had the greatest effect, with a P/N ratio close to 1 indicating no secretion (Fig. 2C, D). An immunofluorescence assay (IFA) confirmed that E proteins accumulated within the cytoplasm of the cells (Fig. S2). This shows the importance of the E-H1 helix in VLP assembly and highlights the critical sites that possibly modulate VLP secretion through the interaction of E-H1 residues facing the lipid bilayer.

**Fig. 2.**
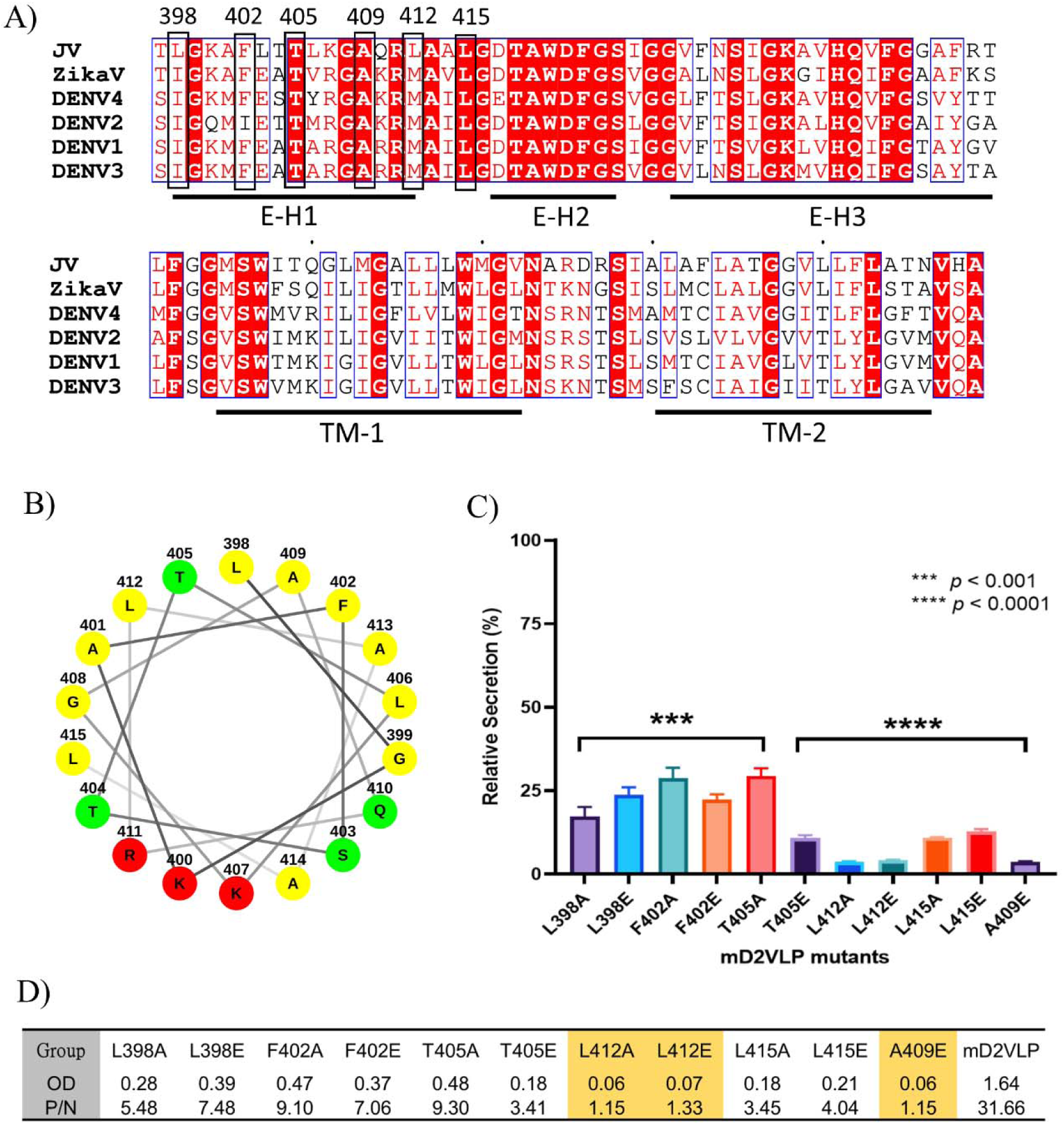
Interaction of E-Helix 1 (E-H1) of the E proteins from different flaviviruses with the head group of the lipid bilayer. **(A)** Amino acid alignment of the transmembrane domain of four serotypes of dengue virus (DENV1-4), Japanese encephalitis virus (JEV) and Zika virus (ZIKV). **(B)** Helical wheel analysis of the transmembrane domain of E-H1 from the JEV E protein. **(C)** The wild-type mD2VLP and site-specific mutated DNA plasmids were transfected into HEK-293T cells by PEI. The respective VLP secretion was determined from the culture supernatant 5 days post transfection by antigen-capture ELISA. The relative secretion level of the respective mutant VLP was calculated as the percentage in comparison with the wild-type mD2VLP. Data are presented as the mean ± standard deviation (SD) of three independent experiments. Statistical analysis was performed using GraphPad Prism, with one-way ANOVA revealing a highly significant difference among the mutant VLPs. **(D)** Summary table of the optical density (OD) and positive-to-negative OD (P/N) ratios for the different mutant VLPs. Notably, certain mutants demonstrated markedly decreased secretion levels with P/N ratio close to 1, compared to others, underscoring the functional impact of these mutations.

To gain mechanistic insights into the effects of the mutations on VLP secretion, we performed a series of 500 ns long AA MD simulations of the dimeric DENV Echi (SH-TM region of JEV) protein complex embedded in biologically relevant lipid membrane models as well as E-H1 mutants. All the systems were embedded in two different membranes (Fig. 3A, B), namely, (i) a phospholipid dominant (PL) lipid membrane (POPC:POPE:POPS) (POPC:phosphatidylcholine (16:0/18:1), POPE:phosphatidylethanolamine (16:0/18:1), and POPS:phosphatidylserine (16:0/18:1)) at a 6:3:1 ratio derived from an insect cell line from the work of Zhang et al.[28] and (ii) a diacylglycerol dominant (DG) lipid membrane consisting of DG:SAPC:FA (DG:diacylglycerol (18:0/20:4), SAPC:phosphatidylcholine (18:0/20:4), and FA:fatty acid (18:1)) at a 56:28:6 ratio (Fig. 3A-3B) derived from the lipidomics in this study. We first calculated the root-mean square deviation (RMSD) and root mean square fluctuations (RMSF) of each E-M complex for Echi and each E-H1 helix mutant to establish the varying effects of point mutations on the overall dynamics (Fig. S3). The RMSD values averaged over the last 100 ns of the simulations showed that the mutations did not have a significant effect on the overall protein complex structure compared to that of the Echi system (Fig. 3C-D, Movie S1). Thus, the mutations do not disrupt the overall protein complex assembly over the timescale of the simulations. However, at the local level, analysis of the per-residue secondary structure and RMSF of the E-H1 helix during the simulation revealed that the F402E and A409E mutants and, to a lesser extent, L412E exhibited partial unfolding around residues 397-413 (Fig. S4 and S5, Movie S2). We next analyzed the interactions of the E-H1 helices with lipids, which showed that Echi exhibited a greater number of lipid contacts than did any of the A or E mutants (Fig. S6A-B and Fig. S7). This was the case for both the PL- and DG-dominant lipid membrane systems. Subsequent calculation of the per-residue membrane insertion depth with respect to the lipid head groups revealed that the E-H1 helices of Echi were buried more deeply into the bilayer than any of the other mutants (Fig. S6C and Fig. S8) for both lipid compositions. Overall, this comparative analysis of Echi and E-H1 mutants provides a rationale for the reduction in secretion efficiency of VLPs observed in *in vitro* experiments, which arises from local loss of E-H1 helix lipid interactions and, in select cases, partial E-H1 unfolding.

**Fig. 3.**
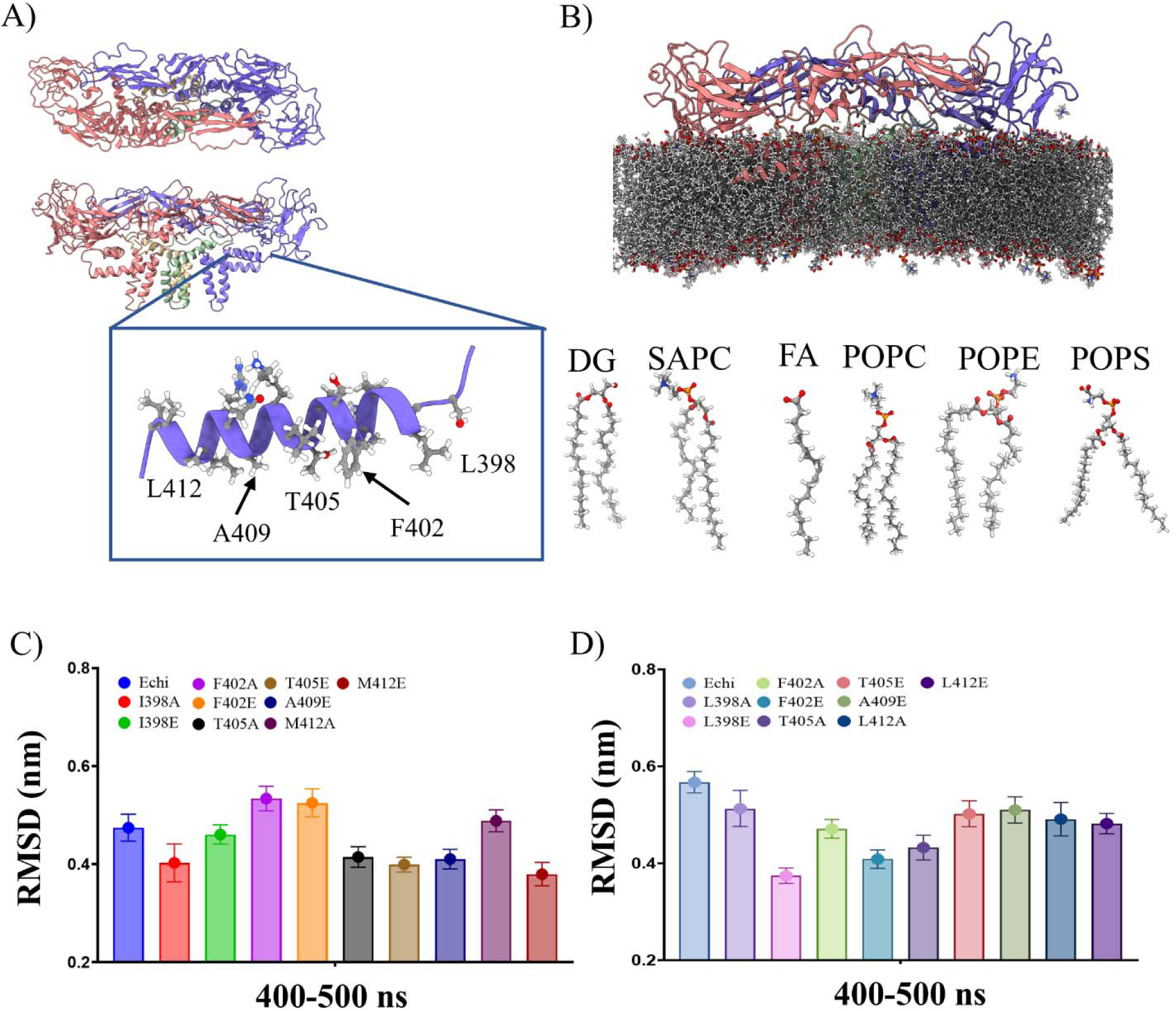
The effect of mutations on the E-H1 helix was assessed using all-atom MD simulations of chimeric E-M protein dimers embedded in two different lipid bilayer compositions. **(A)** A cartoon representation of the E-M protein dimer showing top and side views. The inset shows the magnified E-H1 helix, and the amino acid residues facing the membrane are shown in the liquorice representation. **(B)** AA model of the E-M protein dimer embedded in the lipid membrane. The dominant lipid molecules found in the viral lipid envelope are shown in the liquorice representation. The mean protein backbone RMSD over the final 100 ns of 500 ns long simulation trajectories is shown for Echi and E-H1 helix mutants embedded in **(C)** PL-dominant (POPC:POPE:POPS) and **(D)** DG-dominant (DG:SAPC:FA) lipid membranes.

### E-H1 helix engineering to enhance VLP secretion

In an attempt to increase VLP secretion efficiency, we next aimed to identify mutations in the E-H1 helix that may enhance lipid envelope interactions without drastically perturbing the E protein structure using the Rosetta Membrane Proteins (RosettaMP) module[29] to calculate the difference in free energy between native Echi and each point mutant. A mutational scan was performed at each site of interest, i.e., residues 402, 405 and 409 (sites 398 and 412 are hydrophobic residues), with all amino acids excluding proline across each site. The mutation to leucine had the most favorable (negative) change in free energy, indicating that this mutation may result in a stabilizing effect (Fig. 4A-B). To validate this result, we performed VLP secretion assays with single leucine (F402L, T405L, and A409L), double leucine (F402L/T405L, T405L/A409L and F402L/A409L), and triple leucine (F402L/T405L/A409L) mutations in the Echi protein. Strikingly, both double and triple leucine mutant VLPs produced 2-fold and ∼150-fold greater secretion, respectively, than chimeric Echi and native E protein did (Fig. 4C). Consistent with these findings, 500 ns AAMD simulations of the double or triple leucine mutants of the Echi protein dimer in a DG-dominant lipid bilayer revealed increased helicity of the E-H1 helix, increased E-H1 lipid interactions, and deeper helix insertions into the bilayer (Fig. S9 and Movie S3). This finding supports the notion that critical interactions between the E-H1 helix and the lipid envelope govern the secretion efficiency of DENV VLPs and that this effect may be rationally improved via targeted point mutations. However, further studies are needed to determine whether these findings are generalizable to other DENV serotypes and flaviviruses.

**Fig. 4.**
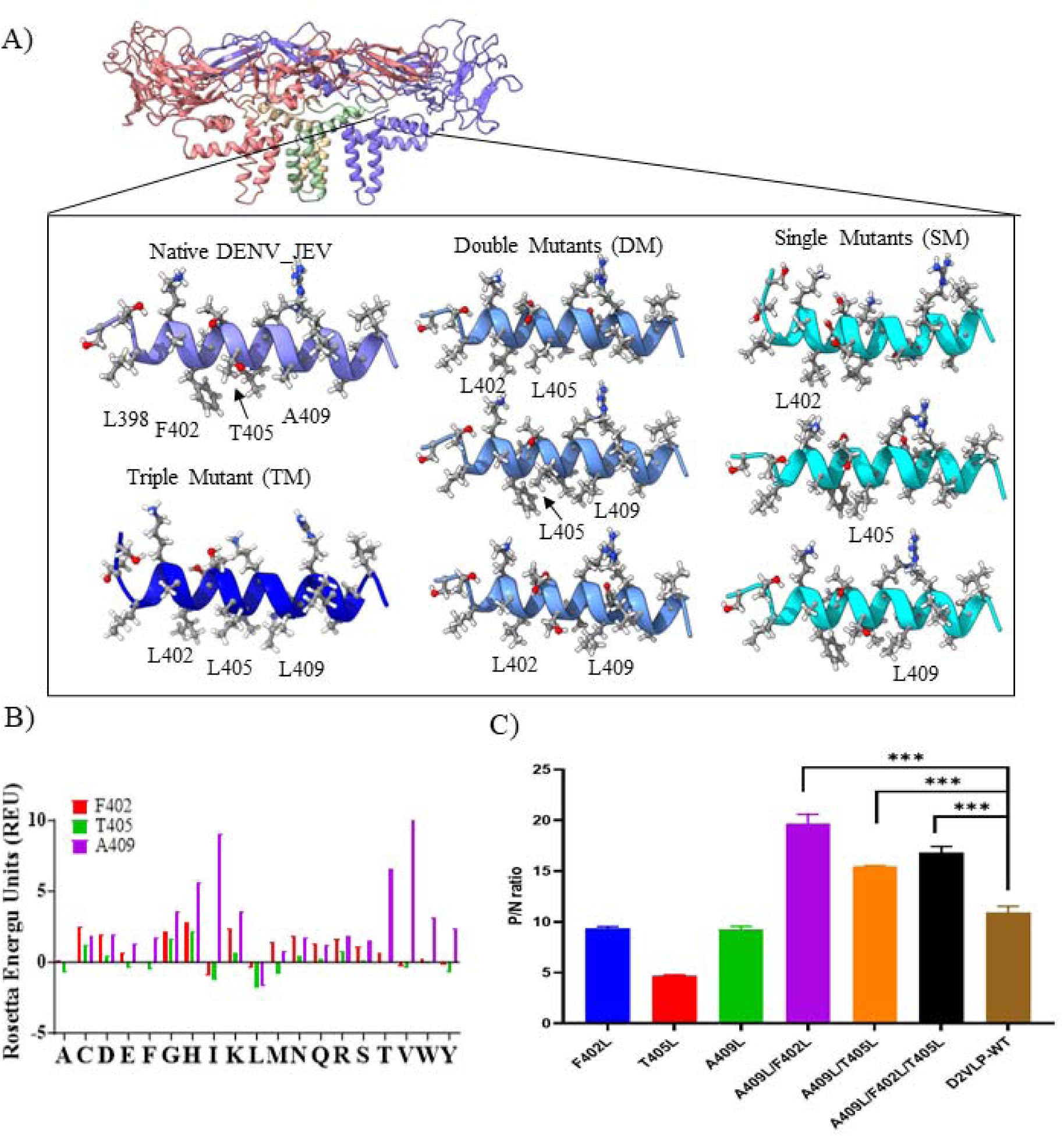
Engineering VLPs with enhanced secretion. **(A)** The E-M protein structure is shown as a cartoon representation, with the inset highlighting stem helix 1 (E-H1) from the native DENV-JEV E protein along with single, double, and triple mutants. **(B) (left)** Plot showing the change in Rosetta energy units (REUs) upon mutation to any other 19 amino acids (proline excluded) along critical E-H1 helix sites, i.e., residues 402, 405, and 409. (**C)** Secretion assay comparing the efficacy of secreted Echi with or without positive mutations, i.e., single/double/triple Leu mutations in the E-H1 helix. Data are presented as the mean ± standard deviation (SD) of three independent experiments. Statistical analysis was performed to determine significant differences between groups, with p < 0.05 indicating statistically significant variation. The results show a marked difference in the secretion levels among the mutant VLPs, with some mutants demonstrating higher secretion levels compared to others.

### EH1-helix mutations enhance homogeneity of VLPs

Further, to assess the effect of secretion enhancing mutations on overall structure of VLPs, we performed cryo-EM on DENV serotype 2 (WT) as well as the highest secreting F402L/A409L double mutant. The purified VLP samples were prepared as described in method section and in our previous work [22, 30]. We observed significant differences in overall size distribution and morphology between the two VLP samples (Fig. 5). Firstly, when looking at cryo-EM micrographs the double mutant shows higher proportion of spherical particles (Fig. 5A, D). We compared 2D-class averages (Fig. 5B and Fig. 5E) and particle size distribution (Fig. 5C and Fig. 5F) from WT and double mutant based on VLPs micrographs. In case of double mutant, a single dominant 2D-class average consisting of 40% of total number of particles was observed. In case of WT each dominant 2D class consisted of 14-23% of total number of particles (Fig. 5B and Fig. 5E). This indicates that secretion enhancing mutations contribute to producing homogenous particles as a result of enhanced protein-lipid envelope interactions. The number of particles in dominant size range i.e., 30-32 nm is ∼46% and ∼33% for double mutant and WT respectively (Fig. 5C and Fig. 5F). This further confirms the homogeneity attained in mutant VLPs. In summary, the wild-type sample displayed significant size and shape variability among the particles, consistent with observations reported by Shen et al [21]. In contrast, the double mutant sample exhibits a single size peak, indicating more uniform particles with clear symmetry, suggesting tightly packed proteins on the particle surface.

**Fig. 5.**
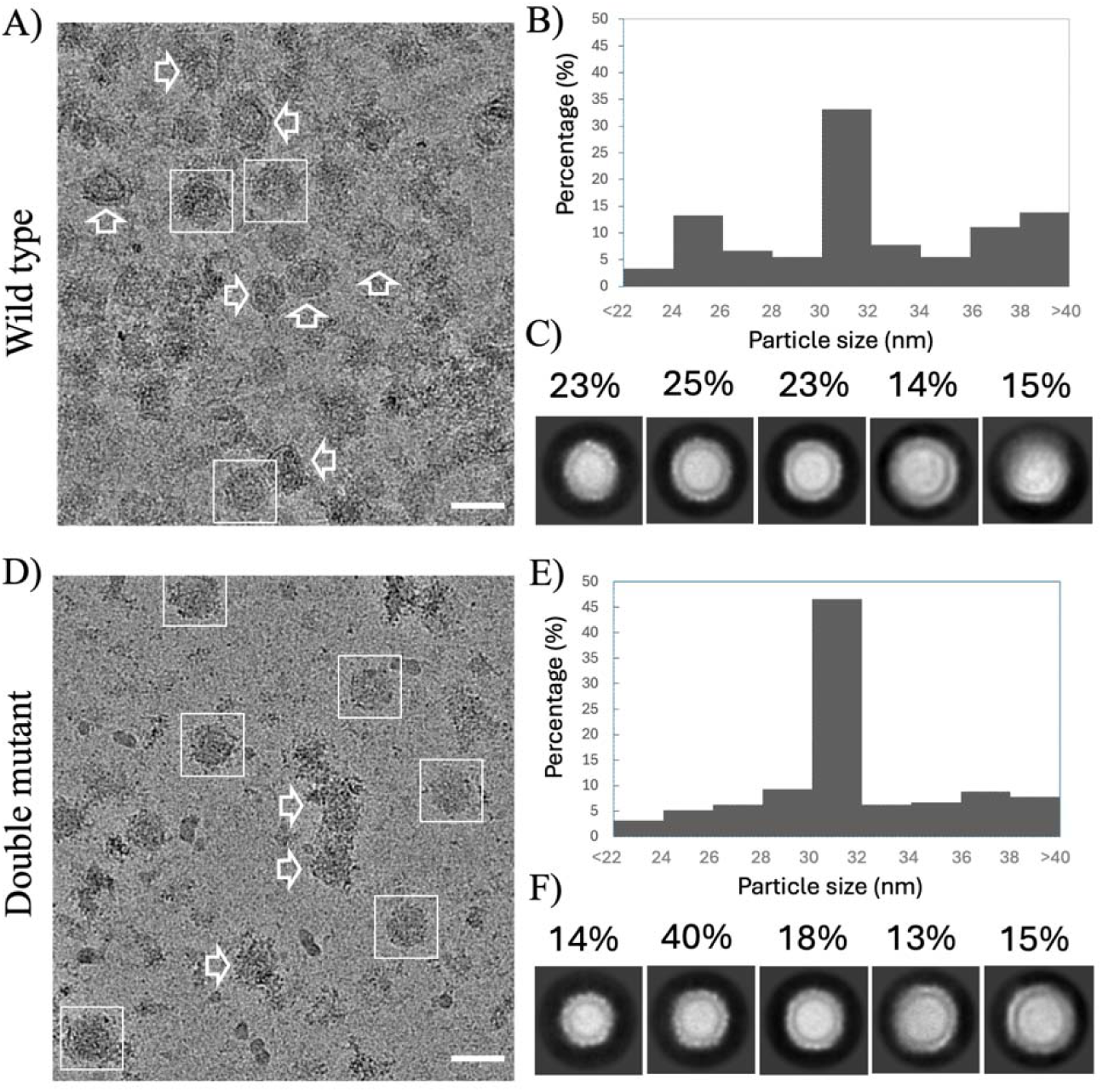
Cryo-EM Analysis of Wild-Type and Double Mutant Mature Dengue Virus-Like Particles. Representative cryo-EM micrographs of wild-type and double mutant mature dengue virus-like particles showing spherical particles (boxed) and irregular or incomplete structures (arrows) are shown in **(A)** and **(D)** respectively. The scale bar corresponds to 50 nm. Size distribution of wild-type and double mutant samples are shown in **(B)** and **(E)**. Selected 2D class averages of double mutant and wild-type mature dengue virus-like particles, obtained through reference-free 2D classification using CryoSPARC v3 are shown in **(D)** and **(F)**. Particles ranging from 24 to 38 nm in size were included in the analysis.

### Coarse-grained (CG) modeling of mature VLPs

Previous studies have shown that antibodies can target epitope sites on the E protein monomer or dimer or across more complex quaternary sites on the flavivirus particle surface[8]. Epitope exposure is a crucial variable among vaccine candidates that governs the magnitude of the host immune response[22]. Therefore, we constructed a fully assembled model of the mature VLP to gain insight into epitope exposure in the context of native protein protein quaternary interactions and the lipid envelope. We used a CG simulation approach to explore the dynamics of the entire VLP over physiologically relevant timescales. A protein shell consisting of 30 E-M dimeric units was constructed by fitting the AA protein structures into the lower-resolution cryo-EM density map of the VLP (EMDB:6926) and embedding into a lipid vesicle with an ∼90 Å radius guided by the corresponding density (see the Materials and Methods section for details). A total of 6 VLP models were built based on two vesicle lipid compositions, i.e., PL-dominant (VLP_PL_) or DG-dominant (VLP_DG_) membranes, each with three alternatives for the number of lipids packed into the vesicle (VLP_PL_-1, VLP_PL_-2, VLP_PL_-3 and VLP_DG_-1, VLP_DG_-2, VLP_DG_-3) (Fig. 6A-5C). Each VLP model was first subjected to a 500 ns equilibration simulation, during which the protein backbone beads were restrained. This allowed lipid molecules to freely rearrange around the E-M protein shell, adopting a stable configuration inside the region of density for the lipid envelope indicated by the low-resolution cryoEM map. In the VLP_PL_ systems, the lipid molecules became uniformly distributed around the protein shell (Fig. 6C). In the VLP_DG_ systems, the spontaneous formation of lipid “bulges” or a local increase in bilayer thickness at specific regions was observed, as shown by calculations of head group density distributions (Fig. 6D). This difference appeared to be due to DG molecules accumulating in the space between the bilayer leaflets of the vesicle, resulting in bulge formation beneath the E-M protein shell. Multiple previous studies have reported the accumulation of neutral lipids such as DG between lipid bilayer leaflets, known as transbilayer activity [31, 32]. Confirming that this difference arose from the presence of DG lipids, simulations of the lipid vesicles in the absence of the protein shell revealed that the dominant PL vesicles retained their spherical shape, whereas the dominant DG vesicles exhibited lipid bulge formation (Fig. S10).

**Fig. 6.**
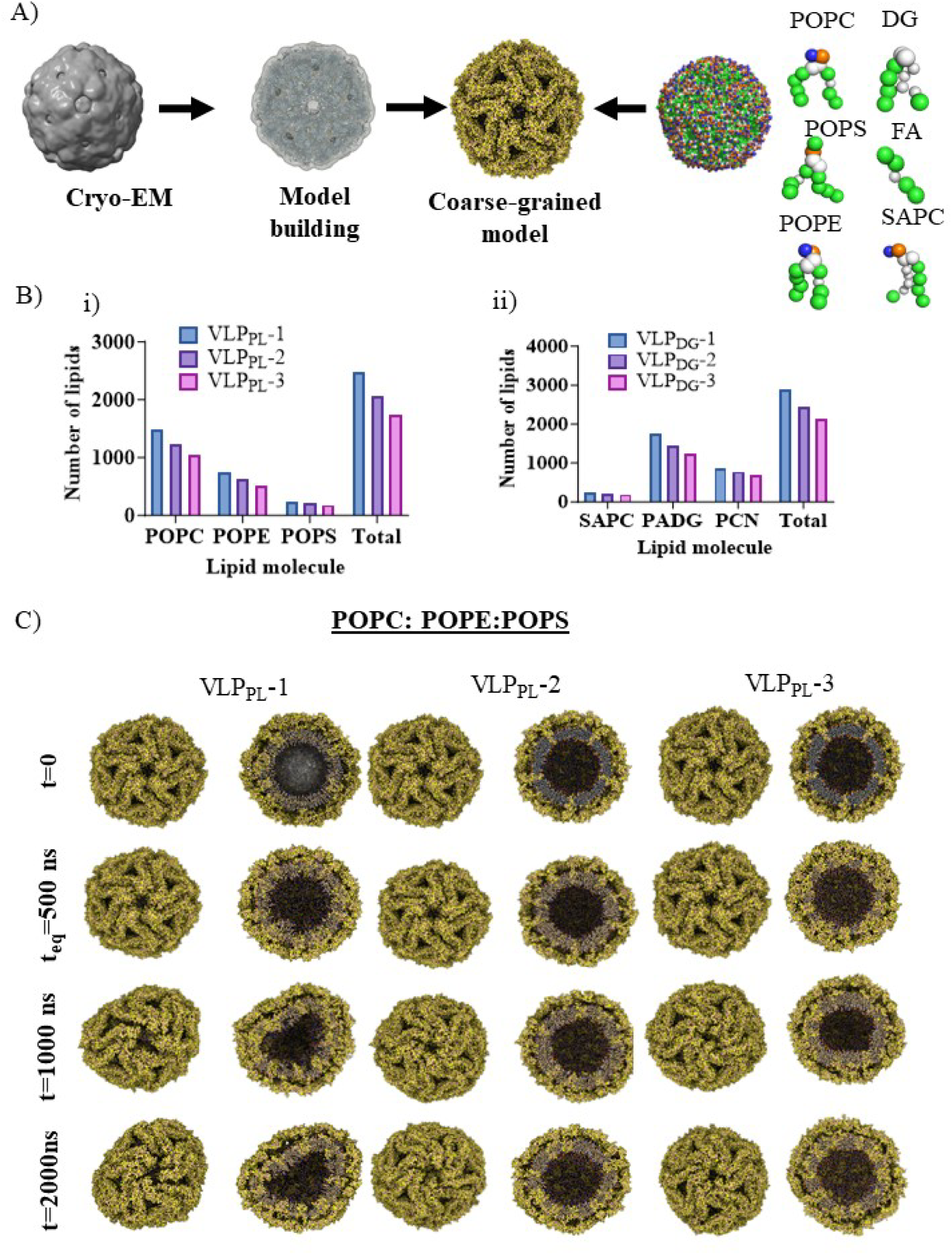
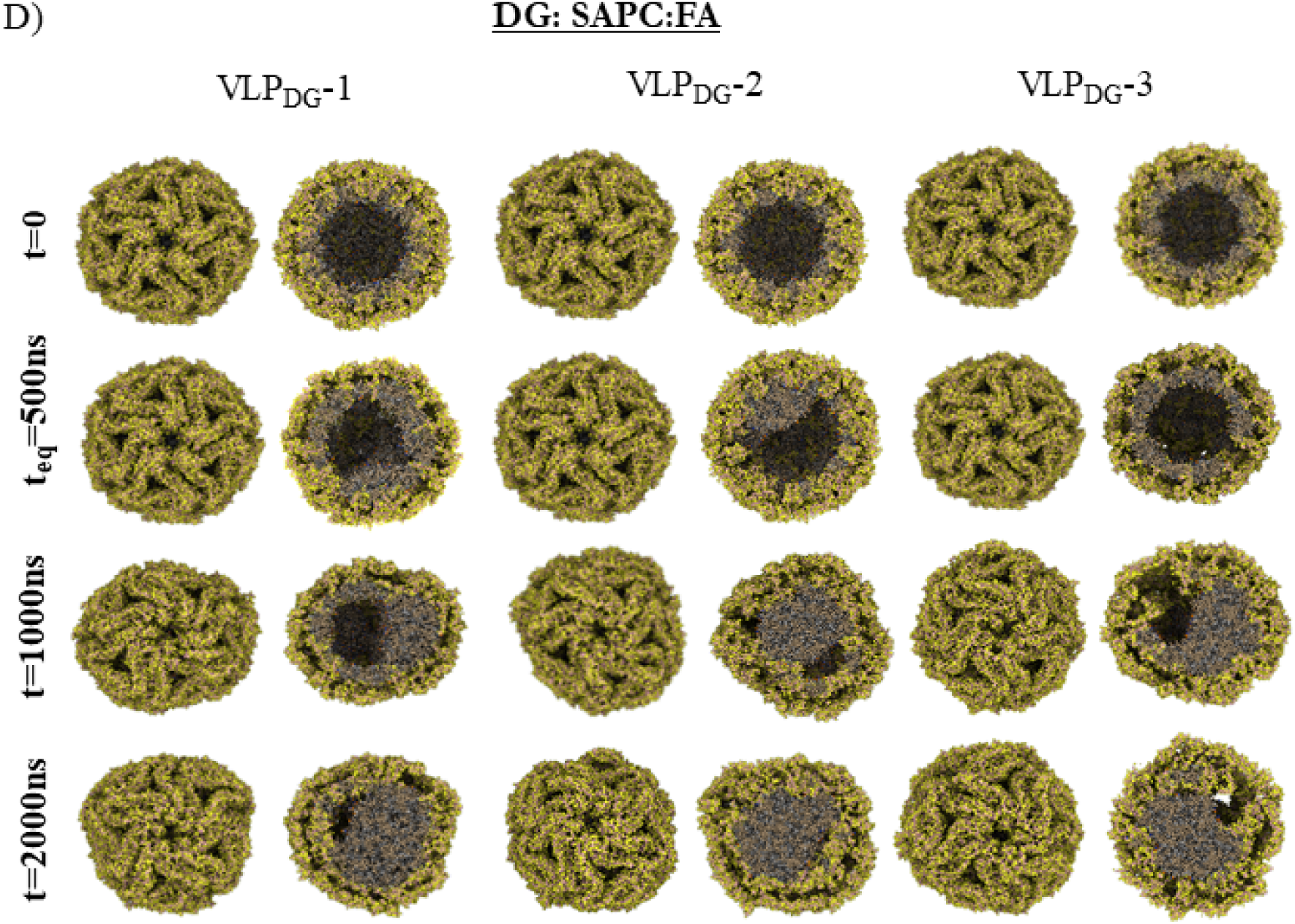
Coarse-grained (CG) simulations of dengue VLPs. **(A)** Workflow of the construction of a CG model of DENV-2 VLPs using an experimentally derived cryo-EM density map (EMD:6926). The composition and number of lipid molecules in each VLP model are shown in **(B)**. Simulation snapshots of VLPs with different lipid compositions are shown for the **(C)** POPC-dominant and **(D)** DAG-dominant VLP models. Snapshots are shown at progressive simulation time points for alternative models involving different amounts of lipids and membrane compositions. Each unrestrained simulation was performed in triplicate. VLP models showing the overall shape and cross-section are shown in bead representations with protein components in yellow and lipid components in gray.

The final equilibrated frame of each VLP model was used to perform 2 µs long unrestrained simulations, which were run in triplicate. Simulation of the VLP_PL_ and VLP_DG_ systems led to distinct effects in terms of VLP morphology (Fig. 6C-D, Fig. S11A). The spherical shape of the VLP_PL_ was dependent on the total lipid concentration (Fig. 6B-D). The VLP_PL_-2 and VLP_PL_-3 systems exhibited a stable spherical shape throughout the simulation time, whereas VLP_PL_-1, which had the highest number of lipid molecules, adopted an ellipsoidal shape, and the structural deviation was greater than that in the simulations with a spherical morphology (Fig. 6C-5D, Movies S4 and S5). The ellipsoidal models are consistent with the particle morphologies previously reported to be present in cryo-EM micrographs, unlike spherical reconstructed structures, due to averaging[22]. This approach helps to rationalize the significant number of nonspherical particles observed in negative stain images or cryo-EM electron micrographs of VLPs. In contrast, in all the VLP_DG_ systems, the lipid bulges that formed during equilibration further expanded, occupying a larger space in the interior of the VLPs underneath the protein shell (Fig. 6D). These data emphasize that the lipid molecules incorporated into VLPs dictate their overall morphology and size.

Our previous work highlighted the presence of a groove at the E protein dimer interface on the mature chimeric DENV VLP, which is absent from the mature DENV virion, indicating that VLPs potentially present unique epitope signatures to prime host immunity[22]. Therefore, we characterized the structural changes in the surface-exposed structural domains of the E protein (DI, DII and DIII). The changes in intra/inter-E protein dimeric contacts as a result of inherent molecular motions of VLPs are masked in a low-resolution structure; therefore, MD simulations allowed us to capture the changes in domain exposure from each VLP model system. To characterize epitope site exposure in the VLPs, we calculated the solvent accessible surface area (SASA) of DI, DII, and DIII (Fig. S11B-D, Table S4). The different VLPs showed distinct propensities for domain exposure, which was in line with the morphological heterogeneity discussed above. Additionally, the SASA values at the starting state (t=0) and the mean SASA values over the simulation time of the domains from each VLP model system were compared (Table S4). Compared with VLP_PL_-3, which had the highest number of lipids, which had the highest exposure of all three domains, indicating that the overpacking of lipid molecules results in the disruption of native contact between E dimers. Consistent with these findings, calculations of the distances between all 30 E-M protein dimers revealed that the interdimer interfaces in VLP_PL_-1 fluctuated the most (Fig. S12A). In contrast, the domain SASAs and interdimer stability were far more uniform in the VLP_DG_ systems, suggesting that the bulging phenomenon in the DG-dominant vesicles may serve to “buffer” the particle size and better maintain the optimal arrangement of the DENV virion’s native E protein shell for antibody generation. Furthermore, calculations of the average distance between DIIIs at each of the 12 5-fold symmetry axes lying diametrically opposite each other showed that DIIIs were clearly distributed in order of particle lipid mass for the VLP_PL_ systems, whereas the VLP_DG_ systems were again more evenly distributed (Fig. S12E-S12F and Fig. S10B). Furthermore, we performed principal component analysis (PCA) on the whole VLP to capture dominant motions. PCA revealed large-scale motion, which supported the morphological changes observed during the simulation (Fig. S12B-S12C). We further investigated the protein lipid interactions between native DENV serotype 2 (VLP^N^-1 and VLP^N^-2) and chimeric (VLP-1 and VLP-2) VLPs using AA (E-M dimers embedded in PL and DG-dominant lipid bilayers) and CG-MD simulations (Fig. S13). Our simulations highlighted the critical differences in the stem-helix-TM (SHTM, residue 395-495) region with lipid interactions in both dimeric all-atom and fully assembled VLP-CG MD simulations (Fig. S13), which contributed to the differences in secretion levels reported previously. Furthermore, a comparison of the sequences of the stem helix transmembrane (SH-TM) regions of native DENV and JEV revealed that the JEV (chimeric) sequence consists of more crucial hydrophobic residues, such as leucine and phenylalanine, which favor prominent lipid interactions (Fig. S14). Overall, our VLP CG simulations reveal the role of E-M protein shell–lipid interactions in influencing morphology and modulating VLP dynamic motion and epitope exposure.

## Discussion

The main challenges of dengue vaccine development are to generate neutralizing antibodies against all four serotypes and morphologically mimic the mature virion with the minimum risk of ADE. VLP, which can be generated spontaneously during the flavivirus infection, has long been explored as a potential vaccine platform. The design of orthoflavivirus VLP needs to consider the VLP assembly process and yield based on different expression systems, which usually produce VLP with different conformation, size and properties [22, 35]. Various approaches in generating orthoflavivirus VLPs, such as DENV, WNV and ZIKA, reveal that these particles assemble into various shapes and sizes ranging from 15–60 Å. The structural characterization of DENV VLPs using conventional methods has been hindered by their inherent instability and complex protein-lipid envelope interactions. In this work, we employed a computational and experimental approach to engineer VLPs with enhanced morphology, homogeneity, and secretion.

Here we focused on characterizing E protein lipid interactions in VLPs to improve secretion properties. Experimental assays and AA MD simulations revealed that the E-H1 helix charge and hydrophobicity significantly influence VLP assembly. By introducing mutations distal to E protein epitopes, we retained the native immunogenic properties of the VLPs while enhancing their secretion. This addresses a critical barrier in large-scale VLP production, where low yields during downstream processing hinder commercial viability. Additionally, coarse-grained (CG) modelling provided insights into epitope exposure and particle morphology in the context of complete VLP models. Consistent with previous studies, we observed significant heterogeneity in VLP size (∼26–36 nm in diameter) and shape (spherical and ellipsoidal) [22, 40, 41].

To investigate this further, we tested two lipid vesicle compositions and modelled lipid molecules within cryo-EM maps. For DG-dominant VLPs, CG simulations showed that lipid bulge formation helps maintain particle diameter regardless of lipid count, capturing the heterogeneity observed in cryo-EM micrographs. For PL-dominant VLPs, the lipid count dictated morphology, with simulations identifying a threshold lipid number required for spherical vesicle formation. Importantly, our CG simulations revealed that nonspherical and heterogeneous VLPs, which constitute the majority of produced VLPs, align with experimental observations. This highlights how fine-tuning protein lipid interactions can reduce heterogeneity and present a diverse array of epitopes to prime host immune responses.

The large-scale production of biologics, including antibodies, viral vaccines, and VLPs, relies on optimizing production efficiency [42, 43]. Industrial strategies such as selecting suitable cell lines, optimizing growth conditions, and refining purification protocols are crucial for enhancing yields and reducing costs [44, 45]. Recombinantly produced non-enveloped VLPs (neVLPs) self-assemble in vitro, unlike enveloped VLPs (eVLPs), which require complex assembly and maturation processes. In the case of DENV VLPs, lipid composition varies between cell lines, emphasizing the importance of selecting lipids with high E protein interaction propensity. Our findings demonstrate that minimal mutations enhancing E-H1 helix insertion and lipid interactions significantly improve VLP secretion which are more homogenous and spherical. Exploring different lipid compositions could further enhance secretion efficiency and produce more homogenous particles.

Furthermore, similar to native infectious viruses, DENV VLPs undergo a maturation process triggered by low pH and host proteases [22]. However, studies have reported inefficiencies in this process, leaving many particles in an immature state [46]. In prior work, we introduced mutations at the furin cleavage site of the prM protein, shifting the equilibrium toward mature particles [22]. Future efforts could focus on optimizing protein-lipid interactions at the immature stage to generate VLPs with desired properties. However, the lack of structural data on immature DENV VLPs makes it a challenging task.

In summary, various approaches have been used to produce orthoflavivirus VLP as the potential vaccine platform; however, very few studies explore its morphology and structure stability. In this study, a computational and experimental integrated approach was developed to engineer VLPs with enhanced morphology, homogeneity, and secretion, which provide the proof-of-concept to expand to VLPs generated from different construct and expression platform if the VLP structure becomes available.

## Materials and methods

### Virus-like particles (VLPs) and mutants

Three different previously constructed and characterized recombinant DNA plasmids expressing prM-E of flavivirus were used to generate VLPs from COS-1 cells (monkey fibroblast kidney tissue, ATCC #CRL-1650), namely, mD2VLP, JEV, and ZIKV. Based on the protocol generated in our laboratory, as previously described[47], the mD2VLP-expressing plasmid pVD2 was used to express prM, 80% of the E protein of DENV-2 (Asian one genotype, strain 16681) and 20% (from the carboxy-terminal end) of the E protein of JEV (Nakayama strain). The plasmid was previously modified at the furin cleavage site of prM to generate mature VLPs. Additionally, the pVD2 plasmid was used for site-directed mutagenesis following the manufacturer’s protocol (Stratagene, La Jolla, CA). The primers used for site-directed mutagenesis are specified in Table S1. Traditional Sanger nucleotide sequencing confirmed that all the plasmids contained no mutations other than those indicated.

Equal quantities (4μg) of plasmids expressing wild-type or mutant mD2VLPs were also transfected into HEK-293T cells, which were subsequently seeded into 24-well plates (human embryonic kidney [ATCC]#CRL-3216) using branched polyethylenimine (PEI) (Sigma Aldrich, Inc., St. Louis, MO, USA) with a PEI:DNA ratio of 1:1. The supernatant after transfection was harvested on a daily basis for ELISA up to day 5. Five days posttransfection, the cells were fixed with 70% acetone for immunofluorescence (IFA) assays.

### VLP secretion level determined by antigen-capture enzyme-linked immunosorbent assay (ELISA)

The secretion of VLP-containing supernatant harvested from the plasmid-transfected cells was tested by ELISA as described previously[22]. Briefly, a 96-well ELISA plate (Thermo, #442404) was coated with in-house-prepared rabbit sera against D2VL, ZIKV-VLP or JEV-VLP at 4°C overnight. Then, 50 μL of each transfected supernatant was added to each well and incubated at 37°C for one hour. The panflavivirus anti-E monoclonal antibody FL0231 (a kind gift from Dr. L.-K. Chen, TzuChi University Hospital, Hualien, Taiwan) and the secondary antibody goat anti-mouse-IgG-HRP were added sequentially for detection. The ELISA results for detection were reported according to the average positive-to-negative (P/N) ratio at 450 nm/630 nm for each VLP sample. The 450 nm and 630 nm absorbance readings represent specific wavelengths of light used to measure the optical density of a solution. While 450 nm is the primary wavelength for the colorimetric signal, the 630 nm is here used as a reference wavelength to measure the background noise and to correct for non-specific absorbance. Positive (P) values for each sample were calculated based on the average OD450 subtracted the average OD630 from the replicates that reacted with positive VLP antigens; whereas negative (N) values were calculated similarly from the replicates that reacted with culture medium control without transfection. The P/N ratio was computed after normalization to the negative control (NC_cell_) obtained from the cell culture supernatant via mock transfection. A P/N value of greater than 2 for a given VLP sample was classified as secretion positive. When **P/N ratio is close to 1**, it suggests that the secretion levels observed in the transfected culture supernatant are almost identical to those of the supernatants without transfection, implying no detectable VLP secreted. A helical wheel representation of JEV E-H1 (peptide 398-415) was generated using the NetWheels web application[48].

### Immunofluorescence assay (IFA)

HEK-293T cells were transfected with plasmids expressing mD2VLP wild-type or mutant mD2VL to evaluate intracellular protein expression by IFA. Five days post transfection, the cells were fixed with 70% acetone at room temperature for 10 minutes. The cells were then blocked with 3% bovine serum albumin (BSA) at 37°C for 1 hour. The same panflavivirus anti-E monoclonal antibody, FL0231, was added to the wells and incubated at 4°C overnight. Finally, the cells were washed three times and incubated for 1 h at 37°C with a mixture of fluorescein isothiocyanate-conjugated goat anti-mouse IgG (1:200; Jackson ImmunoResearch, West Grove, PA) and DAPI (4’,6-diamidino-2-phenylindole dihydrochloride; 1:500; Invitrogen, Molecular Probes, Inc., Eugene, OR) diluted in 1% BSA in 1X PBS. After washing 5 times, the coverslips were mounted on glass slides, and the cells were visualized under a fluorescence microscope (CKX41; Olympus).

### Optimization of lipid extraction methods

Three methods were used for lipid extraction of mD2VLP: chloroform extraction, alcohol extraction, and methyl tertiary butyl ether (MTBE) extraction. These lipid extraction methods were previously used to explore the lipid content in virus particles or VLPs[49–51]. Based on a literature search, the chloroform lipid extraction approach is the most commonly used method for extracting virus particles and is modified from the Bligh and Dyer or Folch lipid extraction method[52]. Considering that the lipid composition of mD2VLP has never been explored before, we applied three different approaches to the same batch of purified mD2VLP and compared the yields of each method. Next, 10 μL of purified mD2VLPs from the HEK-293T-transfected group (mD2VLPs) was subjected to lipid extraction. (1) ***After chloroform extraction,*** 800 μl of 0.1 N HCl:CH_3_OH (1:1) was added to the sample, followed by the addition of 400 μl of chloroform and shaking at 4°C for 1 minute. The mixture was centrifuged at 16,000 ×g for 7 minutes to separate the organic matter from the water phase. The lipid-containing organic layer was collected and dried under a chemical fume hood. (2) For ***EtOH extraction,*** 900 μl of acidic ethanol (0.18 M HCl: ethanol=1:3) was added to the sample, which was subsequently subjected to low-temperature ultrasonic shaking for 3 minutes. Then, the supernatant was collected after centrifugation at 14,000 ×g at 4°C and dried under a vacuum concentrator. (3) Methyl tertiary butyl ether (MTBE) extraction: 225 μl of methanol at 4°C was added to the sample, which was subsequently vortexed for 10 seconds. Then, 750 μl of MTBE at 4°C was added to the sample, which was shaken for 6 minutes. The sample was centrifuged at 14,000 ×g for 2 minutes by adding another 188 μl of ddH_2_O and shaking for 20 seconds. The upper organic layer was collected and dried under a vacuum concentrator. All the dried samples were stored at −20°C and redissolved in 0.1% formic acid buffer before injection for LC MS/MS analysis. The mock-transfected group (293T) was also subjected to lipid extraction as a negative control.

### LC-TOF mass spectrometry and lipidomic analysis

Liquid chromatography (LC) time-of-flight (ToF) mass spectrometry (MS) was performed on a Waters ACQUITY UPLC I-Class system with a Xevo G2XS ToF mass spectrometer operated in positive or negative ion mode. The LC system consisted of a Waters Acquity UPLC system. Lipid molecules were separated on an Acquity UPLC BEH Amide column (150 × 2.1 mm; 1.7 μm) (Waters, Milford, MA, USA). The column was maintained at 45°C at a flow rate of 0.4 mL/min. Mobile phase A consisted of H_2_O with 10 mM ammonium formate and 0.125% formic acid, and mobile phase B consisted of ACN/H_2_O (95:5, v:v) with 10 mM ammonium formate and 0.125% formic acid. A sample volume of 0.5−3 μL was used for the injection. The separation of mobile phase B was conducted with the following gradient: 0 min 100%, 2 min 100%, 7.7 min 70%, 9.5 min 40%, 10.3 min 30%, 12.8 min 100%, and 17 min 100%. The sample temperature was maintained at 10°C. The MS scan range varied from 50 to 1200 m/z. The desolvation nitrogen gas was set to 900 l/h at a temperature of 550°C, and the source temperature was set at 120°C. The capillary voltage and cone voltage were set to +2.5/−2 kV and 25 V, respectively. This chromatographic approach allowed effective separation of the different lipid species[53]. Three replicate analyses were performed for each group, first, a full mass scan was performed at 100-1,000 m/z, and the top 20 signals were selected according to the scan strength (the maximum number of candidate ions to monitor per cycle). Parent ions with a valence of 1 and a signal intensity greater than 30 cps were analyzed by mass spectrometry (MS/MS) sequentially through collision-induced dissociation (CID) and subsequently collected at 100-1,000 m/z. The exclusion time for precursor target ions was set at 6 seconds for data-dependent acquisition analysis (DDA). The parameters used were as follows: IonSpray Voltage Flaoting (ISVF): 5,500 V (ES+), −4,500 V (ES-), ion Source Gas 1 (GS1): 50, ion Source Gas 2 (GS2): 50, Curtain Gas (CUR): 27, and temperature (TEM): 450.

The lipidomic data of each VLP from each extraction method were subsequently collected by searching for m/z ratios against the online database Lipidmaps (http://www.lipidmaps.org). The total sum of the m/z spectra of all lipid categories detected from purified mD2VLPs was calculated, and the lipid group extracted with chloroform had the largest sum of the total spectra obtained in the positive or negative ion scanning mode (**Fig. S1**). Since the lipid extraction approach using chloroform was the most suitable lipid extraction method based on the yield, the chloroform method was used to determine the lipid compositions of the different VLP preparations.

### All atom simulation system setup

E protein dimers from dengue (gene id: KU725663.1) and chimeric dengue (80% DENV, i.e., residues 1-394 plus 20% JEV, i.e., residues 395-495) were modeled using Modeler version 9.10[54, 55] with the PDB templates 3J27[28] and 5WSN[56], respectively. The 100 models generated for each E protein dimer and the structure with the lowest discreet optimized protein energy (DOPE) score and highest percentage of residues with dihedral angles in the allowed region of the Ramachandran plot were chosen for MD simulations. AAMD simulations were performed in GROMACS-2018 using the CHARMM36m force field [57]. Native or chimeric or E-H1 helix mutant E-M protein dimers were embedded into two different lipid membrane systems composed of either POPC:POPE:POPS or DAG:POPC:FA (POPC: 1-palmitoyl-2-oleoyl-glycero-3-phosphatidylcholine (16:0/18:1); POPE: 1-palmitoyl-2-oleoyl-glycero-3-phosphatidylethanolamine (16:0/18:1); POPS: 1-palmitoyl-2-oleoyl-glycero-3-phosphatidylserine (16:0/18:1); DAG: diacylglycerol (18:0/20:4); SAPC: 1-stearol-2-arachidonoyl-phosphatidylcholine (18:0/20:4); and FA: Oleic acid (18:1). Lipid bilayers were built using the CHARMM-GUI Membrane Builder,[58, 59] and ratios of the number of lipid molecules were obtained from the current and previous studies. A 60:30:10 ratio of the number of molecules on the phospholipid dominant membrane (POPC:POPE:POPS) was reported for a native viral membrane [60], and a DG dominant lipid membrane (DG:SAPC:FA) ratio of 56:26:8 was used as per the lipidomic data from the present study. The orientation of the E-M protein dimers with respect to the lipid bilayers was predicted using the Orientation of Proteins in Membranes (OPM) reference server[61]. Dimeric E-M protein lipid bilayer systems were solvated with ∼66,000 TIP3P[62] water molecules in a 16.6×16.6×12.2 nm^3^ box and a physiological salt solution of 150 mM NaCl (Table S2) in addition to neutralizing the overall system charge. Single, double and triple mutations at sites 402, 405, and 409 on the chimeric E protein were modeled using the CHARMM-GUI PDB Reader & Manipulator tool[63]. The E-M protein-lipid membrane mixture was subjected to 5,000 steps of steepest descent minimization. The minimized systems were subjected to multiple steps of equilibration, with position restraints applied to the protein backbone and lipid head groups, and the initial force constants gradually decreased from 4,000 to 0 kJ mol^−1^ nm^−2^ and 2,000 kJ mol^−1^ nm^−2^, respectively. The force constants were gradually reduced over a total of 1.5 ns of equilibration simulation, performed first in the NVT ensemble with a 1 fs time step and then in the NPT ensemble with a 2 fs time step. Integration was performed using the Leap-Frog algorithm. Semi-isotropic Berendsen pressure [64] coupling at 1 bar was used during equilibration, while a temperature of 303.15 K was maintained using the Berendsen thermostat. Each system production simulation was run for 500 ns in GROMACS 2018[65, 66]. The particle mesh Ewald (PME) method[67, 68] was used for long-range electrostatic interactions, with a real space cutoff distance of 1.2 nm, and van der Waals interactions were set to a 1.2 nm cutoff distance with a force-switching function applied at 1 nm. The Nosé Hoover thermostat (303 K)[69] and semi-isotropic Parrinello-Rahman barostat [70](1 bar) with 1 ps and 5 ps coupling constants were used for the production runs, respectively. Trajectory visualization and analysis, including RMSDs, solvent accessibility[71], principal component analysis (PCA)[72], and secondary structure[73], were performed using VMD [74] version 1.9.3 along with GROMACS tools [75].

The Rosetta membrane protein (RosettaMP) module was used to predict the ΔΔG of E_chi_ protein mutants. The protocol described by Alford F Rebecca et al. (2015) was used in the present study[29, 76]. Briefly, E_chi_-M dimer protein structures oriented according to the OPM database were used. The input pdb file was cleaned and renumbered using *clean_pdb.py* (from ROSETTA scripts), and span files were generated using the *span_from_pdb* application in the RosettaMP module. To assess the effect of point mutations on stem helix-1 of the E protein, the MPddG module was used, with a fixed backbone prediction protocol and the franklin2019 scoring function[77, 78]. Side chain conformations were sampled for residues within 8 Å of the mutation position, and the ΔΔG was calculated using Rosetta energy units (REUs) between the mutant and native structures. Mutants with favorable predicted ΔΔG xsvalues (i.e., negative) were subjected to MD as described above, and their secretion efficiency was experimentally tested.

### DENV wild type and double mutant VLP Sample preparation

The mature dengue serotype 2 VLPs (wild-type) and F402/A409 double mutant (mutant) were produced by transfecting HEK293T cells with the recombinant pVD2i-C18S plasmid and the mutant plasmid, using the Lipofectamine® 2000 DNA Transfection Reagent (Thermo Fisher Scientific, USA). The culture supernatants were harvested and subjected to purification via 5-25% sucrose gradient centrifugation in TNE buffer (50 mM Tris-HCl, 100 mM NaCl, 1 mM EDTA) using a Beckman SW-41 Ti swinging-bucket rotor. Fractions with the highest OD values in the ELISA assay were collected, and the VLPs were resuspended in TNE buffer. To further concentrate the samples, Amicon® Ultra-0.5 centrifugal filter units with a 100 kDa cut-off were used.

### Cryo-EM data collection and image analyses

Fresh samples were immediately applied to glow-discharged Quantifoil grids (Quantifoil GmbH, Germany) for cryo-EM grid preparation. Excess liquid was blotted off, and the grids were rapidly vitrified in liquid nitrogen-cooled liquid ethane using a Vitrobot Mark IV (Thermo Fisher Scientific). Cryo-EM images were acquired at a magnification of 81,000x and an accelerating voltage of 300 kV, using a FEI Titan Krios transmission electron microscope equipped with a K3 detector (super-resolution mode). The pixel size at the specimen level was 1.061 Å/pixel.Image processing and 2D classification were conducted using CryoSPARC v3 [79]. A total of 7,204 and 7,373 particles from the wild-type and mutant samples, respectively, were selected for heterogeneity analysis. The particles, ranging in size from 15 to 50 nm, were included in the analysis process.

### Coarse-grained simulation setup

ChimeraX[80, 81] was used to construct a fully assembled atomistic model of the VLPs with 60 E-M proteins fitted into the cryo-EM density map of mature DENV-2 VLPs (EMDB:6926)[22]. The fully assembled VLP structure consists of 30 E-M protein dimers embedded in a lipid vesicle[38]. The AA coordinates of the E-M proteins assembled in the VLP structure were converted to a Martini3 CG model[82] using the *martinize.py* script available in the vermouth python module. To retain the native fold, an elastic network (EN) was introduced within each E and M protein monomer. The EN was applied between pairs of beads within a cutoff range of 0.5 to 0.9 nm using a force constant of 1.000 kJ mol^−1^ nm^−2^. The EN was applied only within each domain (DI, DII and DIII) of the E protein ectodomain, while the secondary structure of the E and M protein TM regions was maintained using standard angle and dihedral potentials. The CG E-M protein shell of the VLP was embedded into a lipid vesicle with a radius of ∼90 Å (as per the approximate density of lipids observed in the cryo-EM map) by superimposing the centers of mass of the protein shell and lipid vesicle. Lipid vesicles were generated using the CHARMM-GUI Martini marker [83] with different lipid compositions guided by experimentally identified compositions. The protocol for embedding E-M proteins into lipid vesicles has been described elsewhere[38]. After embedding the E-M protein shell into the respective lipid vesicle, lipid molecules with coordinates overlapping with the protein beads were removed. Multiple initial CG VLP models were generated with different numbers of lipids by removing lipid molecules at 0.1 nm, 0.15 nm and 0.2 nm from one another, corresponding to the VLP_PL/DG_-1, VLP_PL/DG_-2 and VLP_PL/DG_-3 systems, respectively. Each VLP model was centered in a cubic box and solvated with MARTINI water along with solvated sodium and chloride beads corresponding to a 150 mM NaCl concentration.

All the systems were subjected to energy minimization of ≤50,000 steps using the steepest descent algorithm. All the simulations were performed using GROMACS version 2018 [65, 66]. In all the simulations, bond lengths were constrained using the LINCS algorithm, and equations of motion were integrated using the Leapfrog algorithm and a 10 fs time step [84]. Pressure and temperature were maintained at 1 bar (isotropic) and 310 K, respectively, using a velocity rescaling thermostat[85] and Berendsen barostat[64], respectively. Six sets of equilibration steps in the NVT and NPT ensembles were performed by gradually lowering the restraints applied to the protein and lipid beads while progressively increasing the time step. The final NPT equilibration step involved position restraints applied to protein beads for only 500 ns to allow equilibration of lipid molecule positions. Unrestrained production runs of 2,000 ns were run in triplicate for each VLP CG (Table S3). Furthermore, we performed principal component analysis (PCA) on the protein backbone beads of the VLPs for each VLP system from the concatenated triplicate simulation trajectory of 6 µs.

## Acknowledgments

The computational work for this article was fully performed on resources of the National Supercomputing Centre, Singapore (https://www.nscc.sg). The cryo-EM experiments were performed at the Academia Sinica Cryo-EM Facility (ASCEM). This work also used ASGC (Academia Sinica Grid-computing Center) Distributed Cloud resources, which is supported by Academia Sinica.

## Funding

A*STAR AME Young Individual Research Grant (YIRG) number A2084c0160. The National Research Foundation Competitive Research Programme (NRF-CRP27-2021-0003). Bioinformatics Institute (A*STAR) core funds. National Science and Technology Council (112-2320-B-006 -022 -MY3) in Taiwan.

## Author contributions

Conceptualization: JKM, DYZ

Methodology: JKM, PJB, SRW, DYZ

Investigation: VRP, FCC, GWC

Supervision: JKM, DYZ, PJB

Writing—original draft: VRP, JKM, PJB, SRW, DYZ

## Conflict of interests

The findings reported in this manuscript are protected under [Patent Application No. PCT/SG2024/050717] and are the subject of a patent application filed by Agency for Science, Technology and Research (A*STAR) in Singapore. Authors declare no conflict of interest.

## Data and materials availability

The data supporting this article have been included as part of the Supporting Information.

## Notes

### Competing Interest Statement

The authors have declared no competing interest.

