## Supporting Information for "An Integrative Approach to Rational Engineering of Dengue Virus-Like Particles"

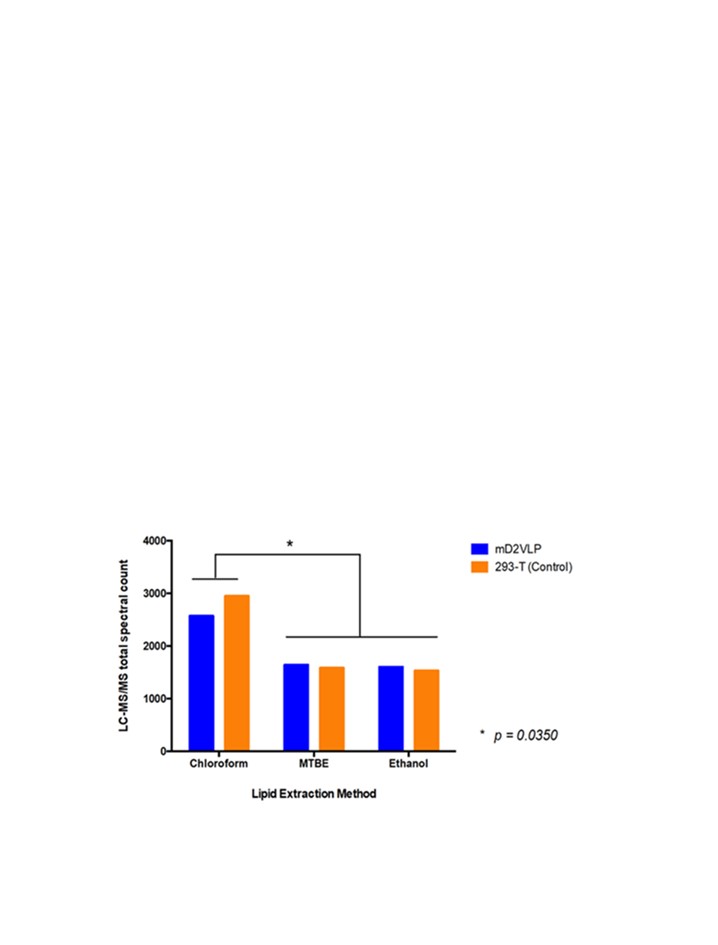

**Fig. S1. Comparison of the extraction efficiency by three different methods of virus-like particles**.

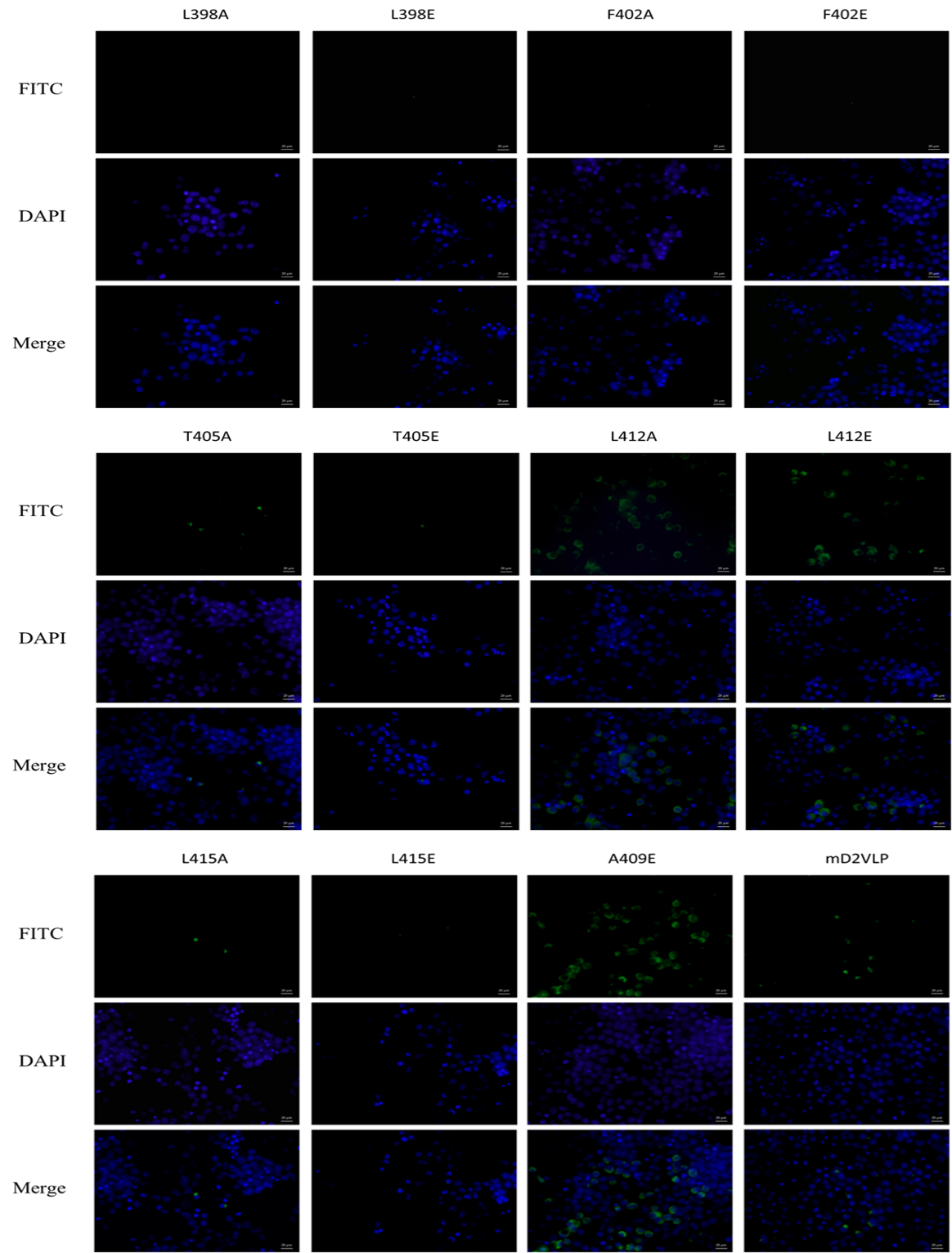

**Fig. S2. Transfection of mD2VLP and the respective SDM plasmid DNA into HEK-293T cells by PEI confirmed the efficiency with IFA.** We performed site-directed mutagenesis (SDM) in E-H1 region and confirmed the plasmid DNA sequences by traditional Sanger sequencing. Based on the best transfection condition, we prepared the plasmid DNA and PEI mixture under a 1:30 ratio and mixed well. Subsequently, we added the master mix into the HEK-293 cells and checked the VLP secretion level after 7 days post-transfection. Meanwhile, the cells were fixed 5 days post-transfection and stained with the primary antibody-D2-MHIAF and the secondary antibody-Goat anti-Mouse-IgG-FITC and DAPI. Using a fluorescence microscope, we could observe the accumulation of green fluorescence in the A409E, L412A, and L412E groups. Those groups have also been shown to have lowered secretion levels in ELISA experiments.

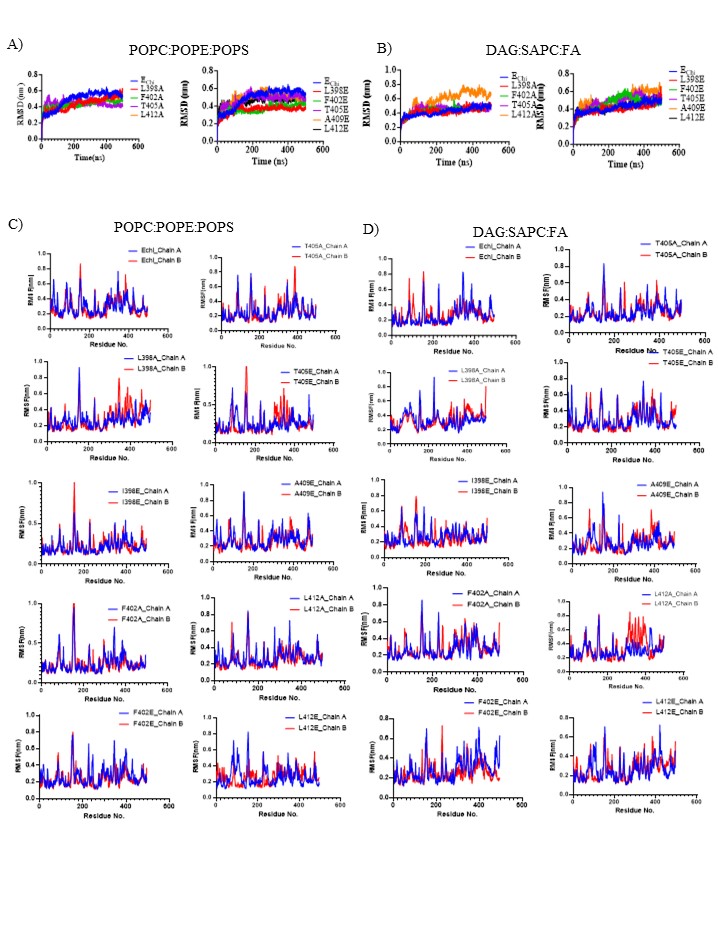

**Fig. S3. Protein dynamics from AA MD simulations of dimeric E-M protein and E-H1 mutants. (A-B)** Plots showing the RMSD of backbone atoms of E protein in Echi and E-H1 mutant systems in PL and DG dominant lipid membranes. (**C-D)** Average RMSF values of Echi protein and respective E-H1 helix mutants embedded in PL and DG dominant lipid membrane systems from 500 ns long AA MD trajectory.

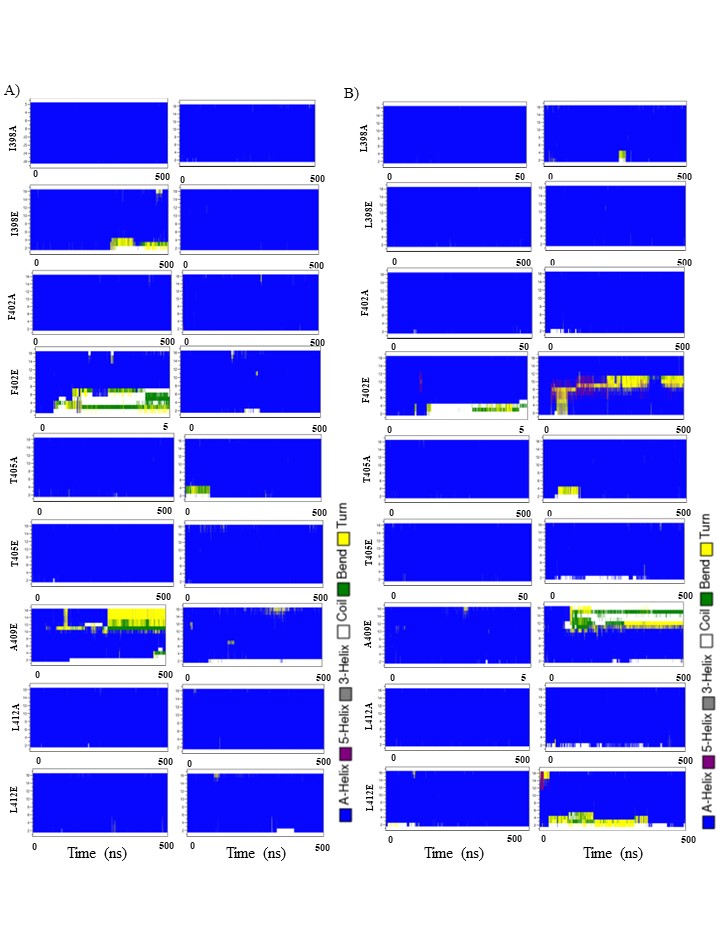

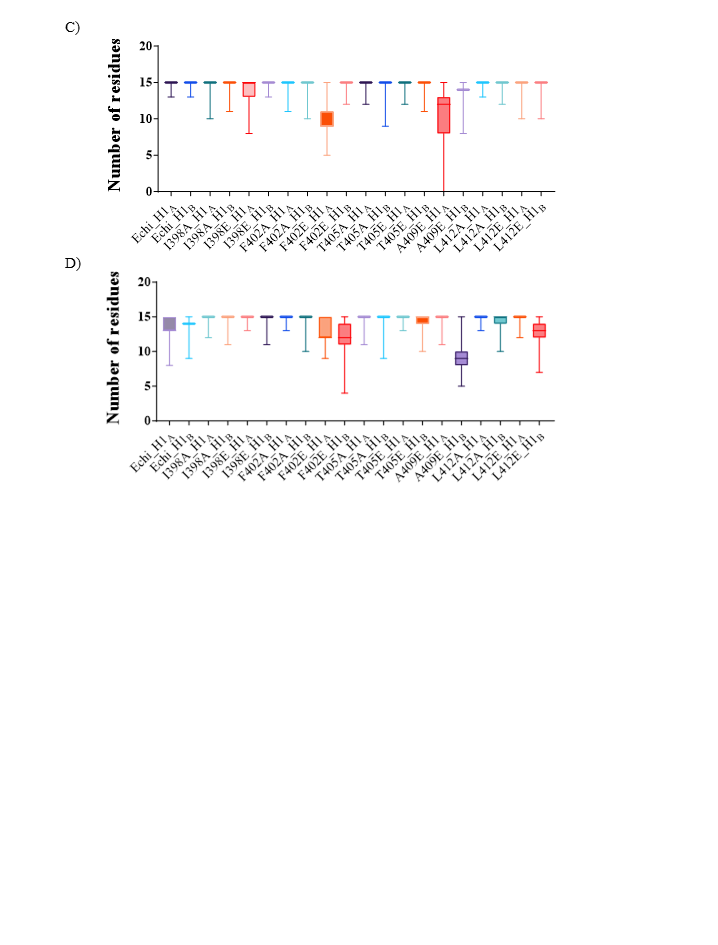

**Fig. S4.** **Secondary structural changes of Ala and Glu E-H1 mutants from AA MD simulations.** Heat maps show secondary structural stability of E-H1 helices (residues 397-413) from 500 ns simulations of Echi-M dimers and its E-H1 mutants embedded in **(A)** PL and **(B)** DG dominant lipid membranes. The secondary structure is colour coded as per key. Box plots show number of amino acids from E-H1 that adopt α-helical structure during the 500 ns simulations Echi-M dimers and its E-H1 mutants embedded in **(C)** PL and **(D)** DG dominant lipid membranes.

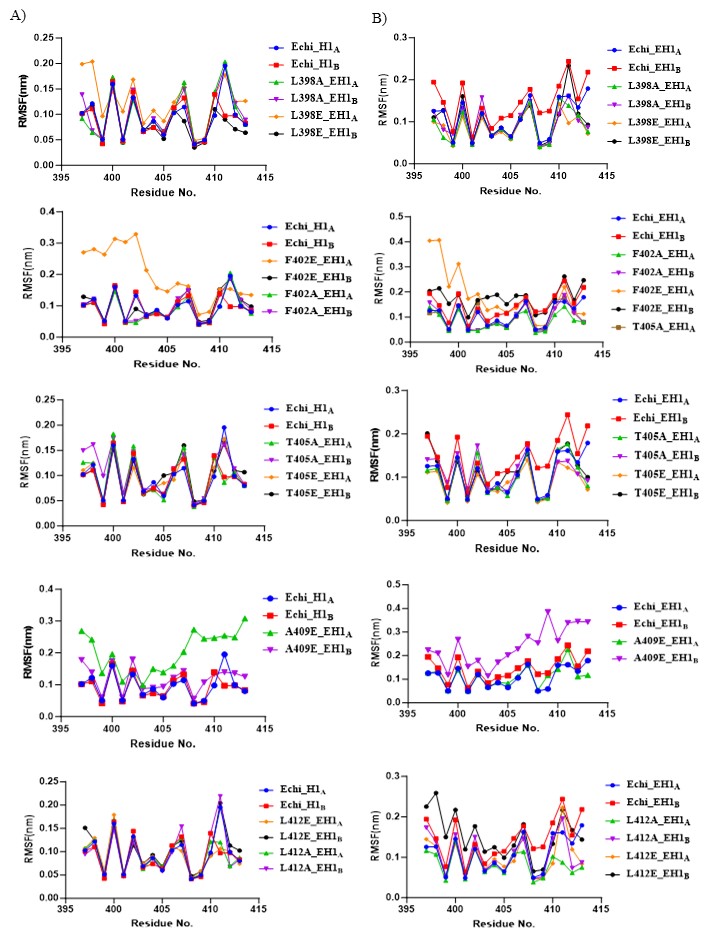

**Fig. S5. RMSF values of E-H1 helix protein from AA simulations.** Average per residue RMSF values of E-H1 helix amino acid residues alone comparing Echi protein and E-H1 helix mutants embedded in **(A)** PL and **(B)** DG dominant lipid membrane systems from 500 ns long AA MD trajectory.

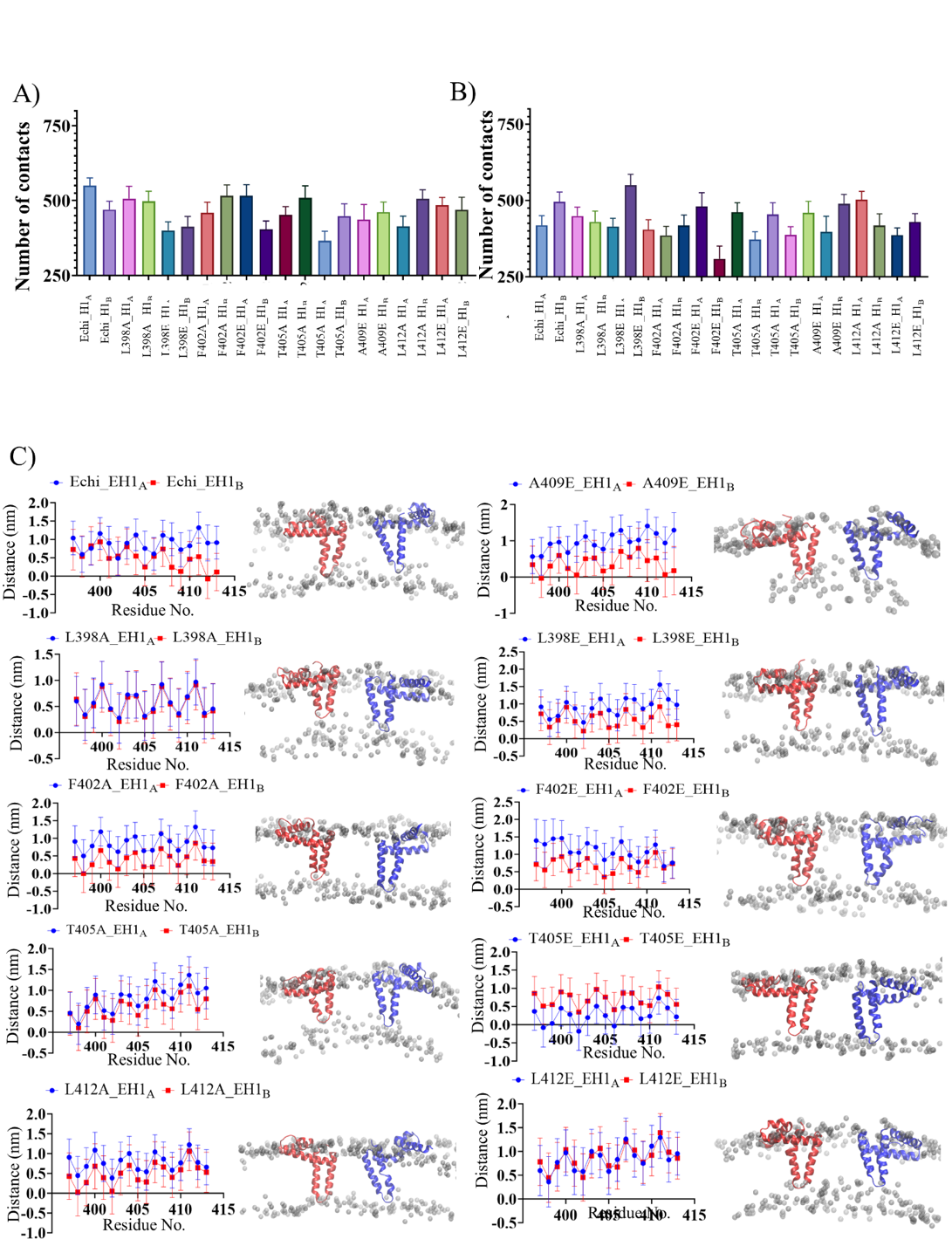

**Fig. S6.** **Per-residue insertion of depth of E-H1 helix within lipid bilayers**. **(A-B)** E-H1 - lipid membrane interactions averaged over the final 100 ns of 500 ns trajectories of Echi and E-H1 helix mutants embedded in either PL dominant or DG dominant lipid membranes respectively. **(C)** E-H1 helix insertion depth with respect to lipid headgroups of E-M dimer in DG dominant membrane simulation. Each point represents the average over the last 200 ns. Alongside each plot, a snapshot of final coordinates of the E-H1 helix is shown. M protein of Echi protein is shown as transparent cartoon and lipid reference plane containing atoms shown as transparent beads. E-H1 helices are shown in solid colors depicting the orientation with respect to the membrane. Error bars were calculated as standard deviation over the final 200 ns of the trajectory.

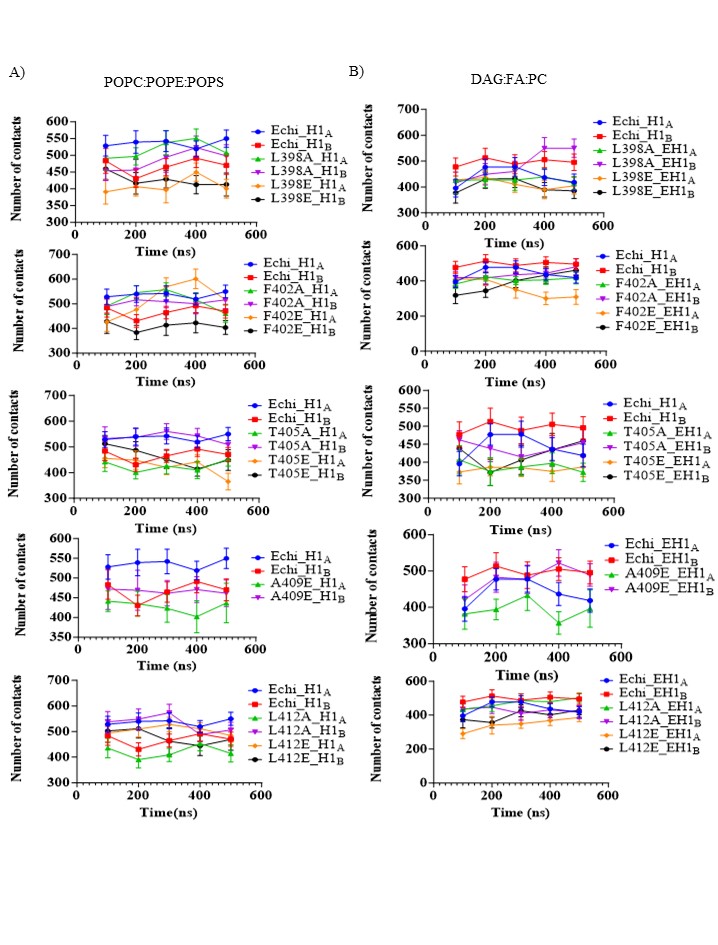

**Fig. S7. Time averaged E-H1 helix-lipid contacts extracted from AA simulations.** Data is shown for **(A)** PL and **(B)** DG dominant lipid bilayers. The error bars correspond to standard deviations from given 100 ns time frame.

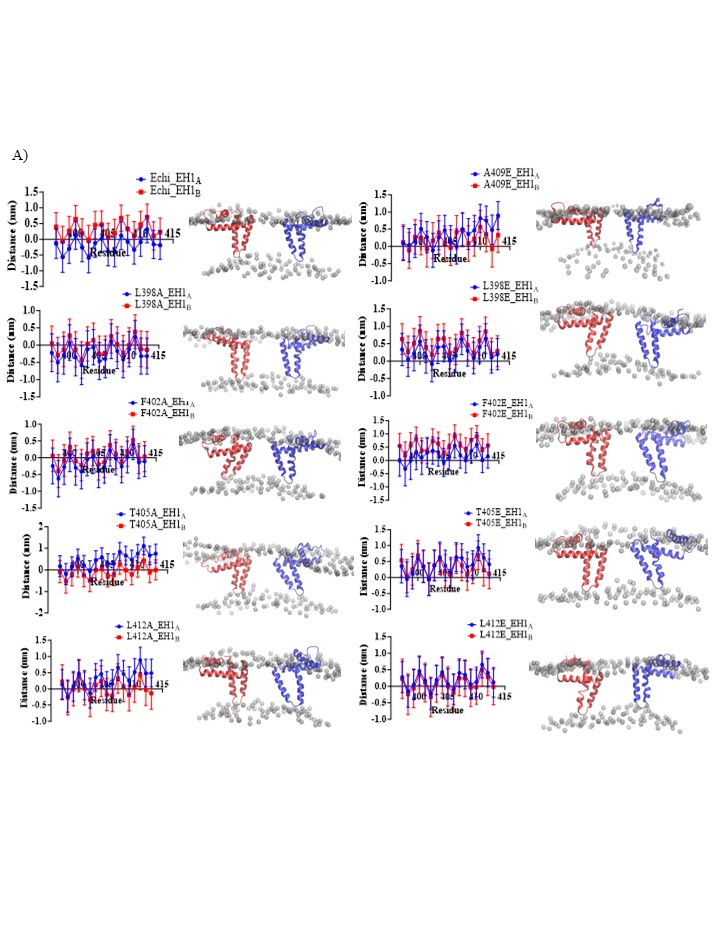
 **Fig. S8. E-H1 helix insertion depth of PL dominant E-M dimer during AA simulation.** Each point represents the average over last 200 ns of the insertion depth with respect to lipid headgroups. Lipid headgroups are defined by a plane corresponding to lipid phosphates in case of PL-dominant lipid bilayer and O31 groups in case of DG-dominant lipid bilayer. Alongside each plot, a snapshot of final coordinates of E-H1 is shown. Echi and M proteins are represented as transparent cartoon and lipid reference plane containing atoms are shown as transparent beads. E-H1 helices are shown in solid colours depicting the orientation with respect to membrane. Error bars were calculated as standard deviation over final 200 ns of the trajectory.

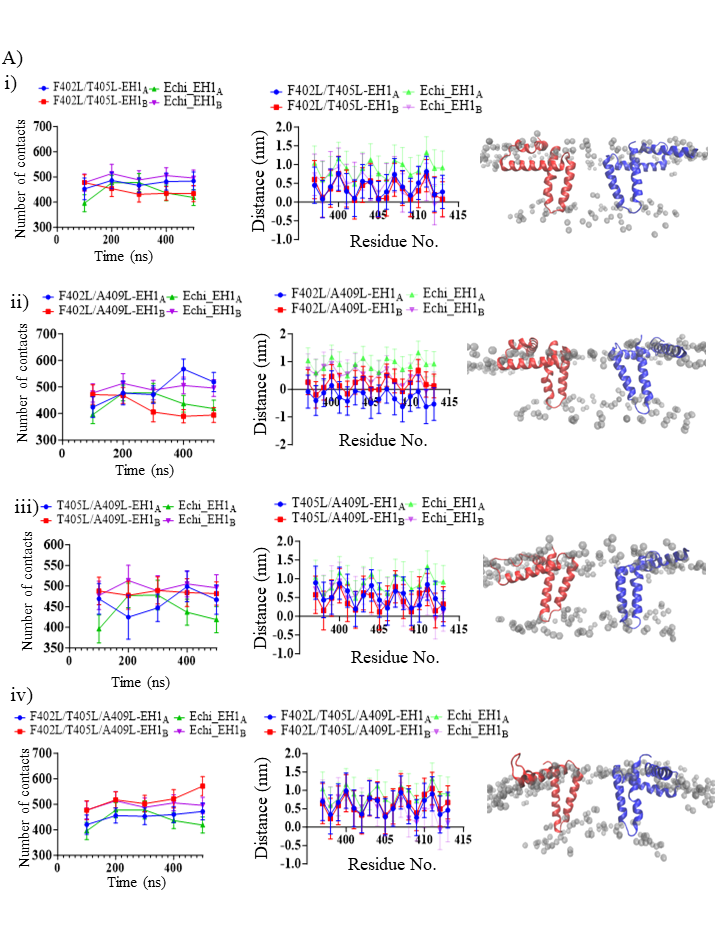

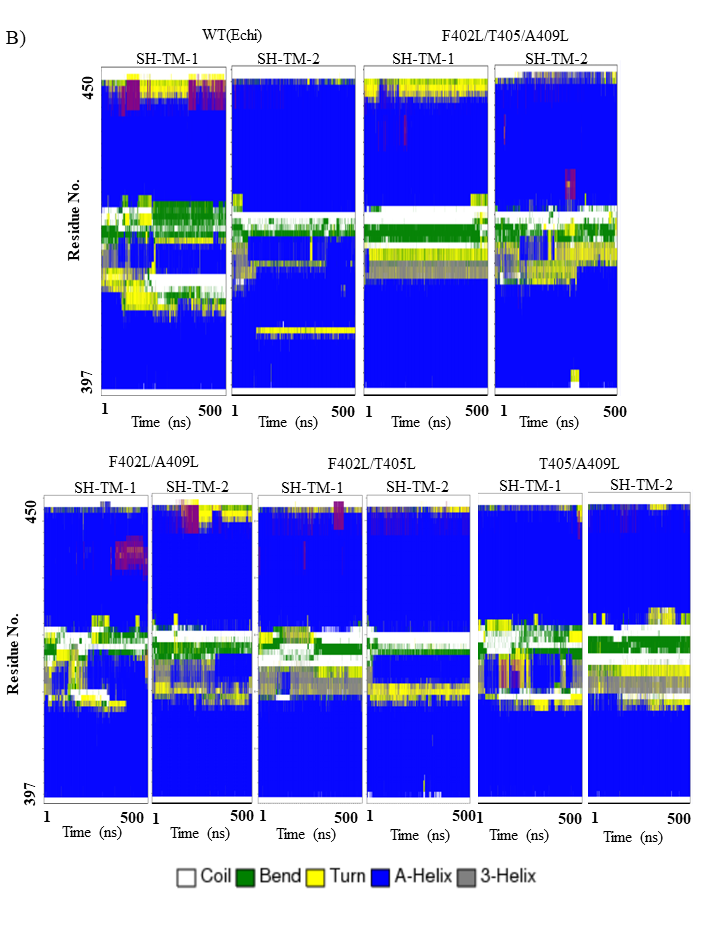

**Fig. S9. Secondary structural changes during AA MD simulations of VLP secretion enhancing E-H1 mutants.** **(A)** EH1-helix: lipid interactions and E-H1 helix insertion into lipid bilayer measured from 500ns simulations performed on E_chi_-M dimer embedded in DG:SAPC:FA membrane. Block average of 100ns is reported for E-H1 helix: lipid interactions and per-residue insertion depth of last 200ns reported for E-H1 helix insertion.

**(B)** Heat map showing secondary structure of stem helix region (E-H1 and E-H2) (residues 397-450) from double and triple E-H1 mutants during 500 ns simulations of dimeric E protein embedded in DG dominant membrane. The secondary structures are colour coded as per key.

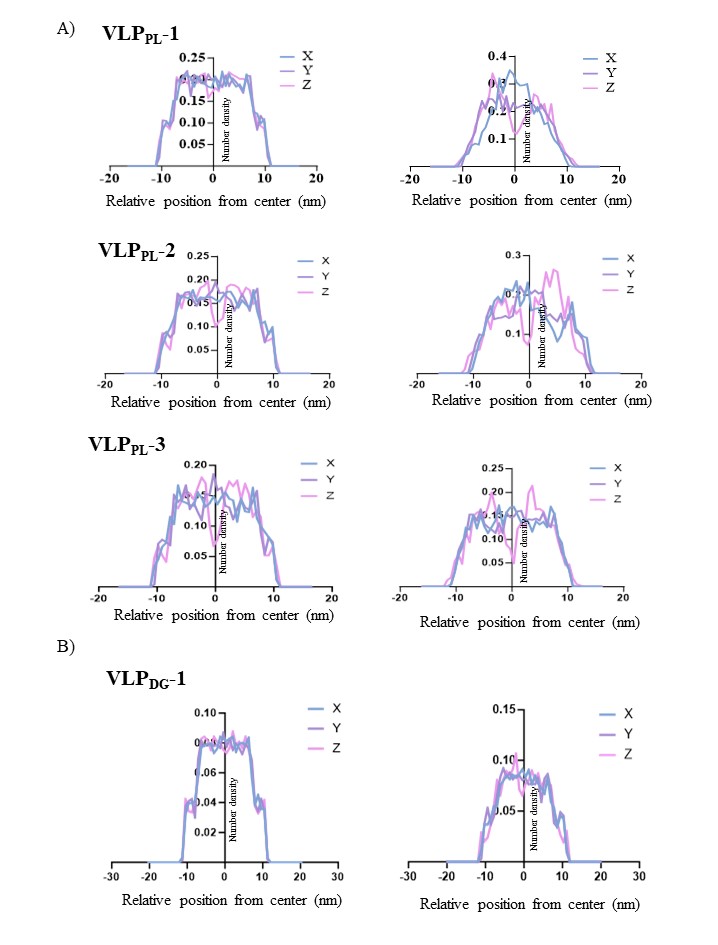

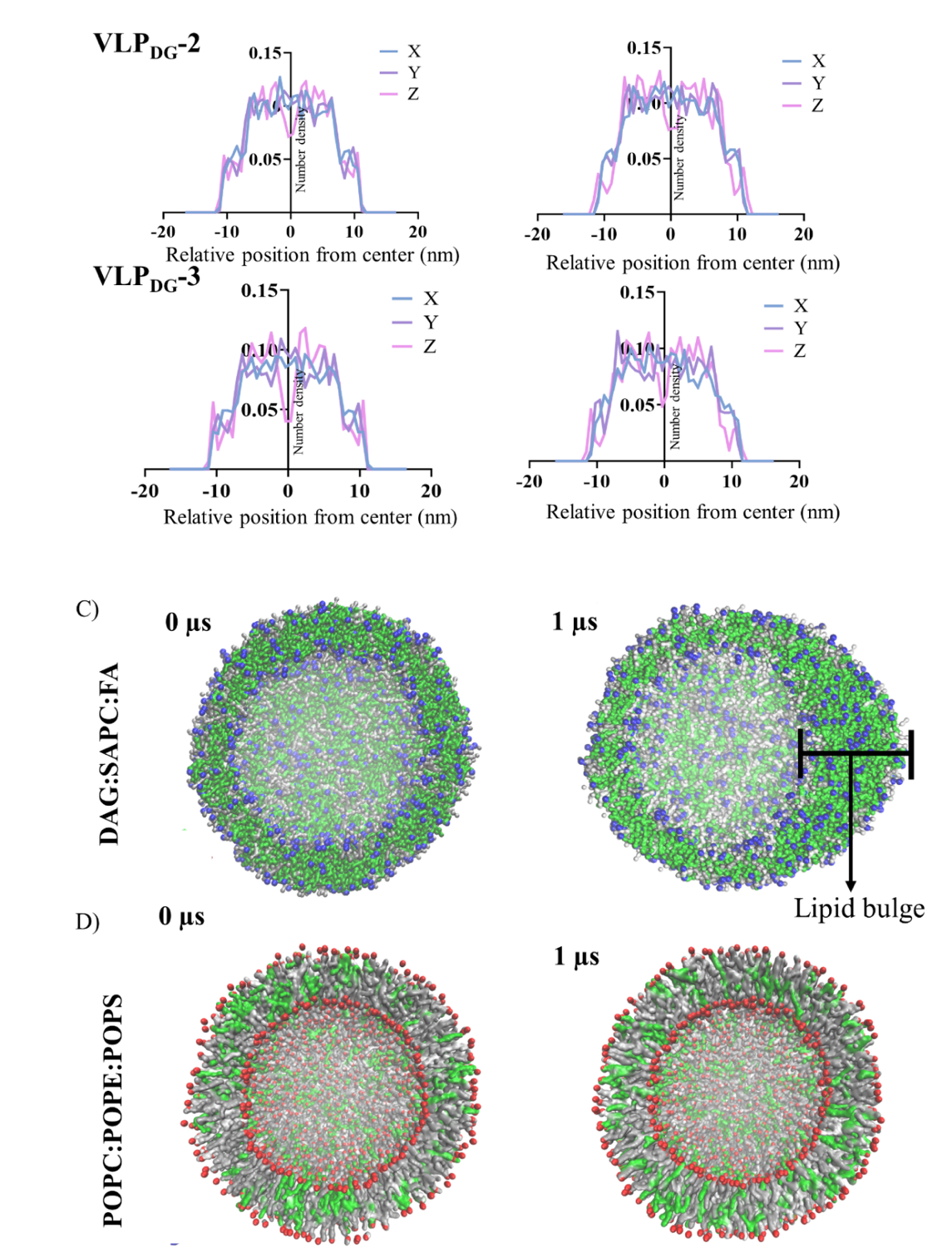

**Fig. S10. MD simulations of Lipid envelope alone reveal DG dependent lipid bulge formation (A)** Phosphate (PO4 bead) head group density distribution along X, Y and Z coordinate from VLP CG-model equilibration simulation. Initial and final corresponds to starting (0 ns) and final (500 ns) configuration of lipid vesicle in VLP. (**B)** GL1 group head group density distribution along X, Y and Z coordinates from VLP CG-model equilibration simulation. Initial and final corresponds to starting (0 ns) and final (500 ns) configuration of lipid vesicle in VLP. Lipid vesicle only simulations of **(C)** DG and (**D)** PL dominant lipid systems showing starting and final structures.

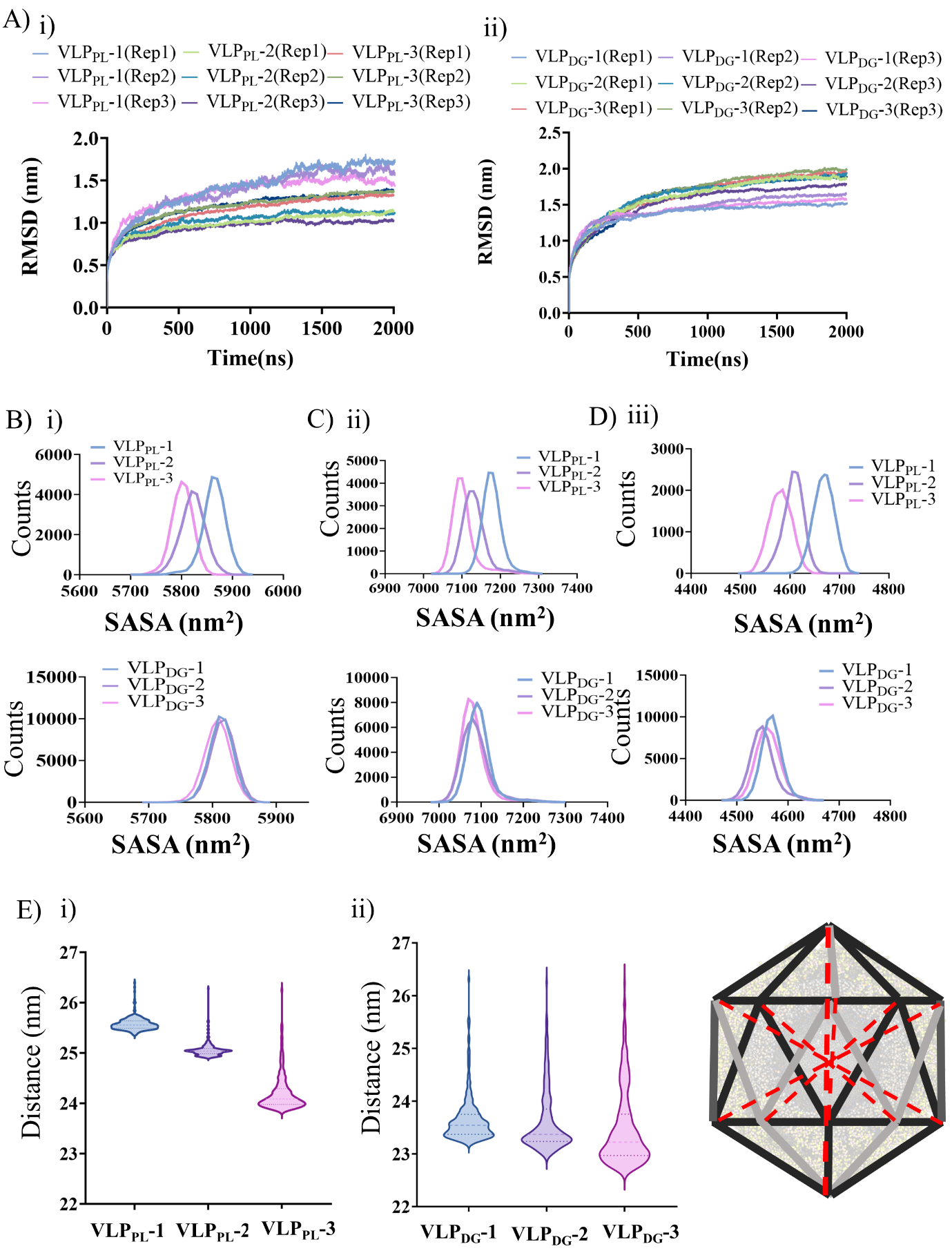

`

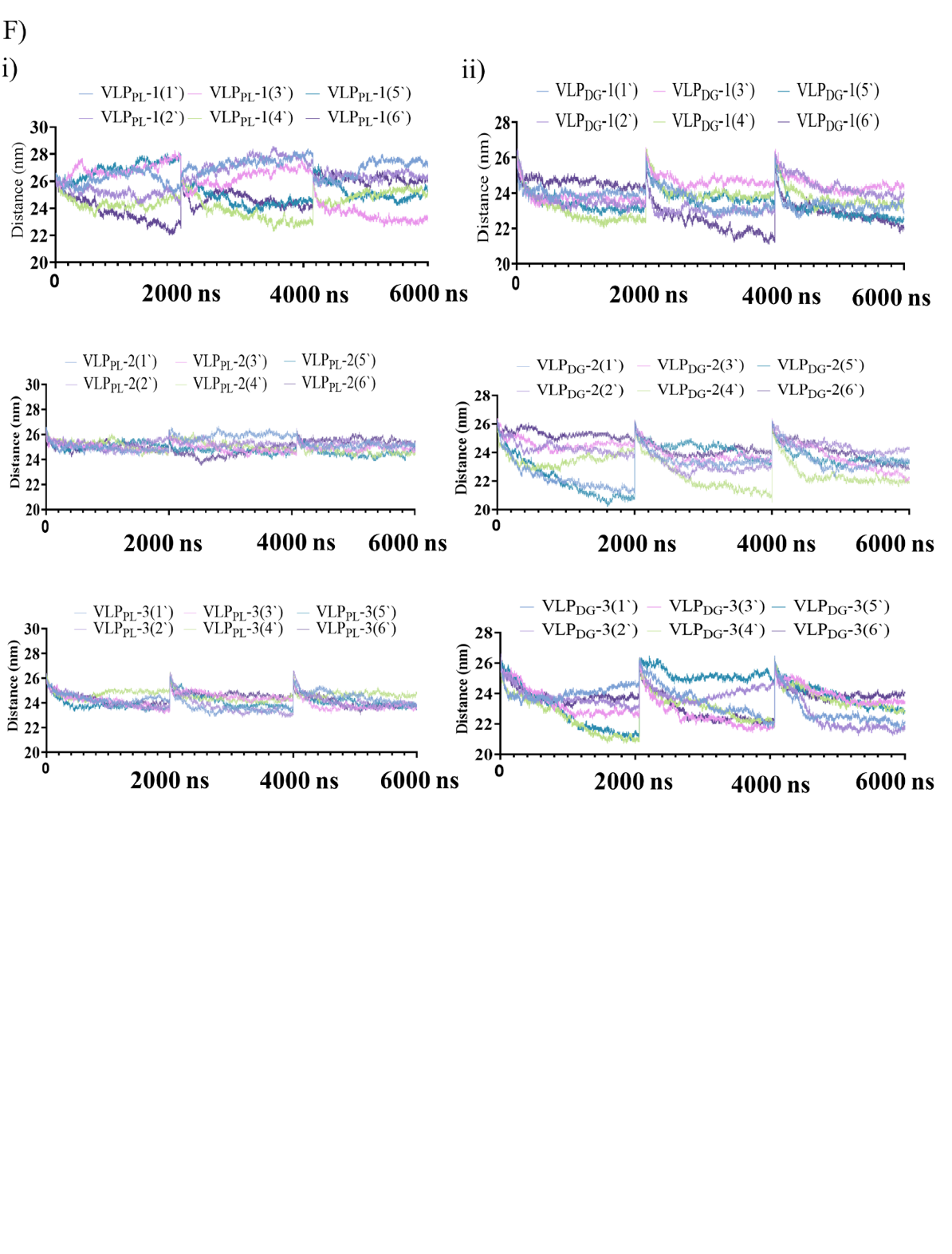

**Fig. S11.** **Lipid composition dependent morphology and dynamics of VLPs. A)** RMSD of VLP backbone beads with respect to the initial structure is shown along the 2000 ns of the unrestrained simulations of three replicates of each model from PL (i) and DG (ii) dominant lipid VLP**.** **(B-D)** Distribution plots of solvent accessible surface area (SASA) of domains (A) DI, (B) DII, and (C) DIII from POPC (top) and DG (bottom) dominant VLP models showing the variation with respect to lipid number. The distribution plots were generated by averaging SASA values measured from 3 independent 2000 ns long simulation replicas of each VLP model . **(E)** Violin plot showing the distribution of average distances between 12 diametrically opposite 5-fold symmetric axes calculated at every 1 ns from a combined triplicate e trajectory of 6000 ns from each VLP model in (i) PL and (ii) DG lipid dominant vesicle systems. Schematic depicting the distances measured between the diametrically opposite symmetric axes in an icosahedral particle plotted in **(E)**. **(F)** Plots showing the distances between diametrically opposite 12 x 5-fold symmetric axes calculated at every 1 ns from a cumulative trajectory of 6000 ns from each VLP model system simulations in PL (left) and DG (right) dominant vesicle systems

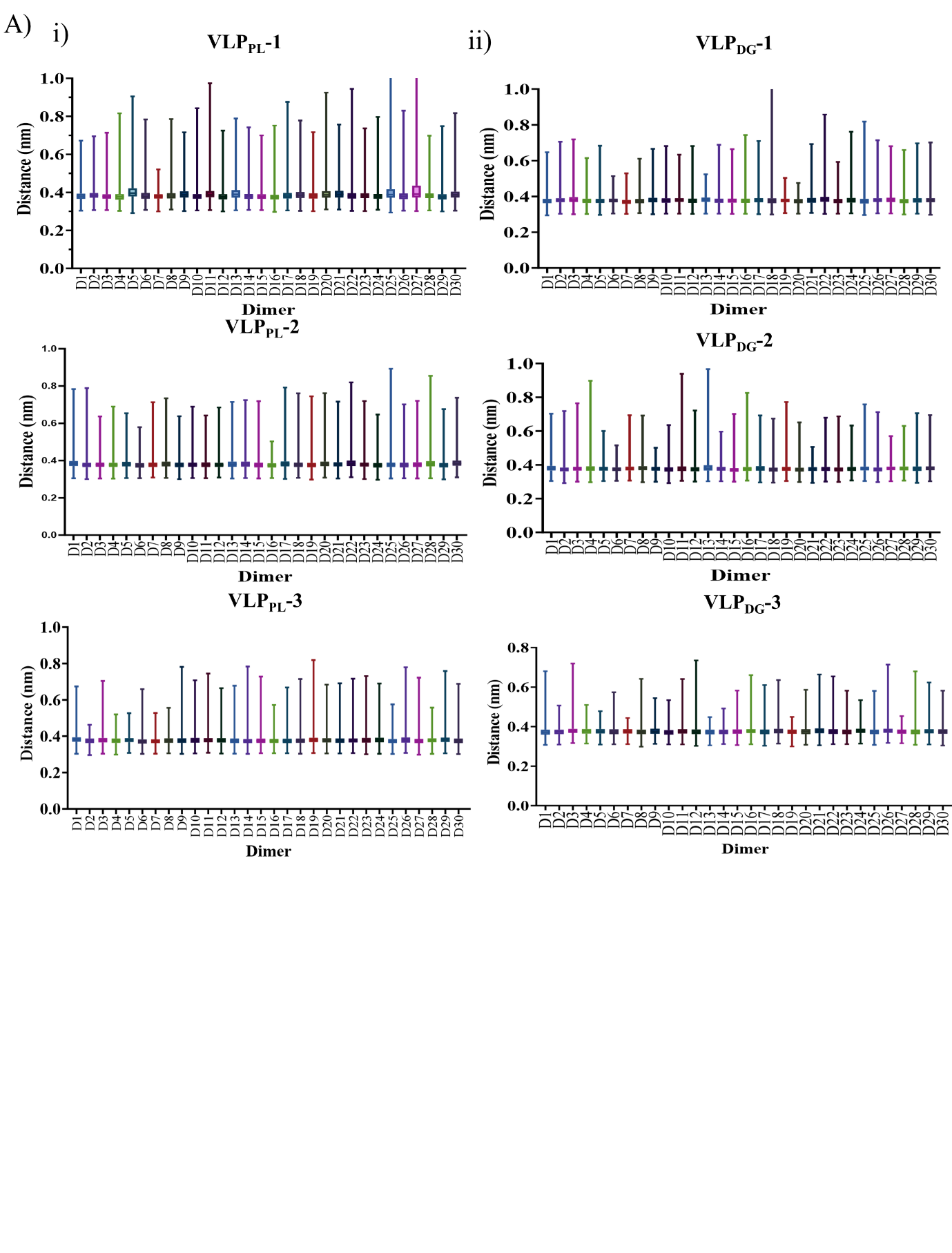

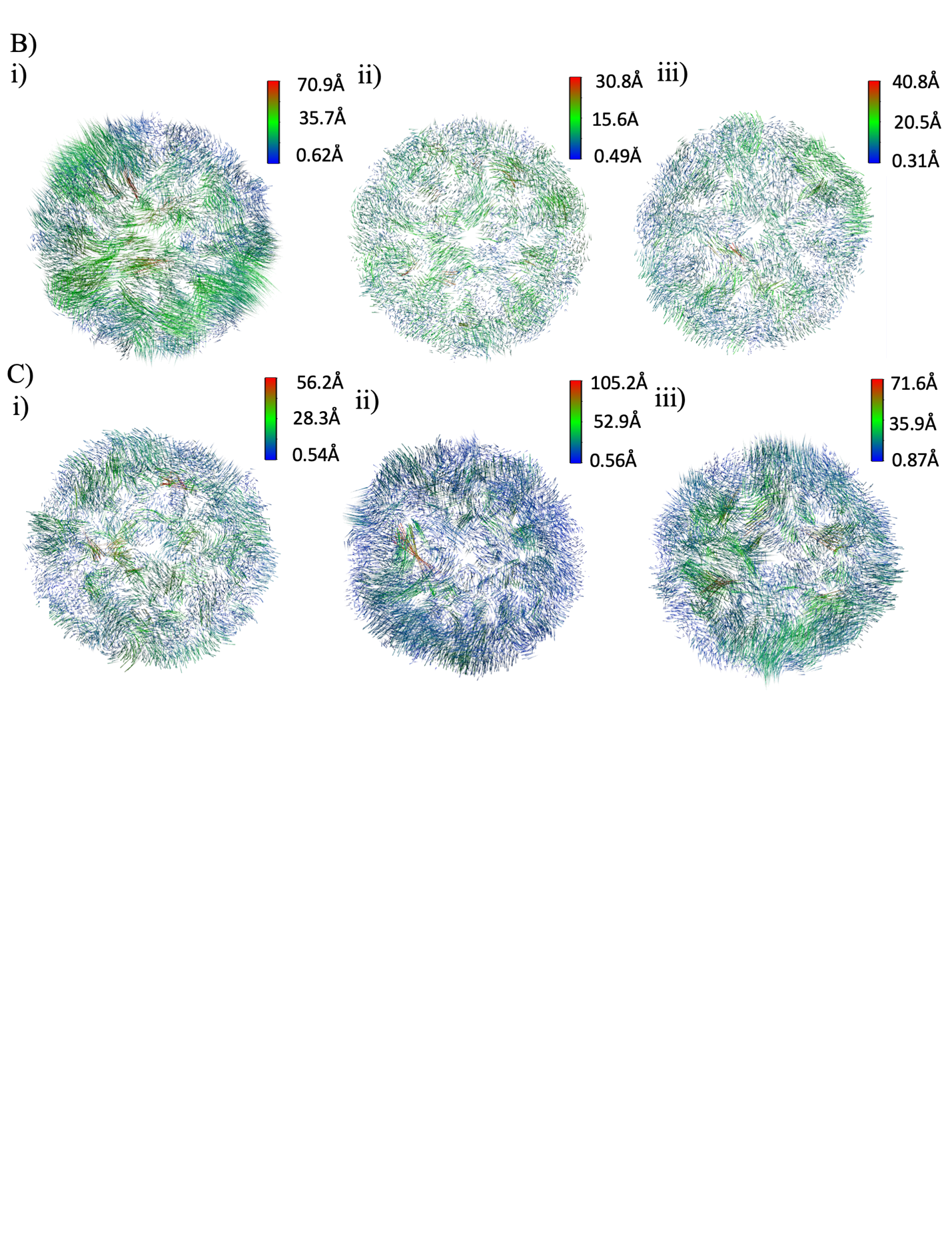

**Fig. S12. CG MD simulations reveal lipid composition dependent morphological changes in VLPs (A)** Minimum distances  between monomers of each dimeric unit of CG-VLP-lipid vesicles models from PL and DG dominant lipid systems of combined triplicate simulations of 6 µs. The dots represent the mean value of distance and error represent minimum and maximum distance. **(B-C)** First principal component mode (PC1) capturing the coordinated motions of each E protein subunit. PCA was performed on every third protein back bone bead of each VLP trajectory from PL-dominant (D, VLP_PL_-1 (i),VLP_PL_-2(ii), VLP_PL_-3 (iii)) and DG dominant (E, VLP_DG_-1 (i),VLP_DG_-2(ii), VLP_DG_-3 (iii)) VLP systems. PC1 accounts for of ~45-50% variation in the data.

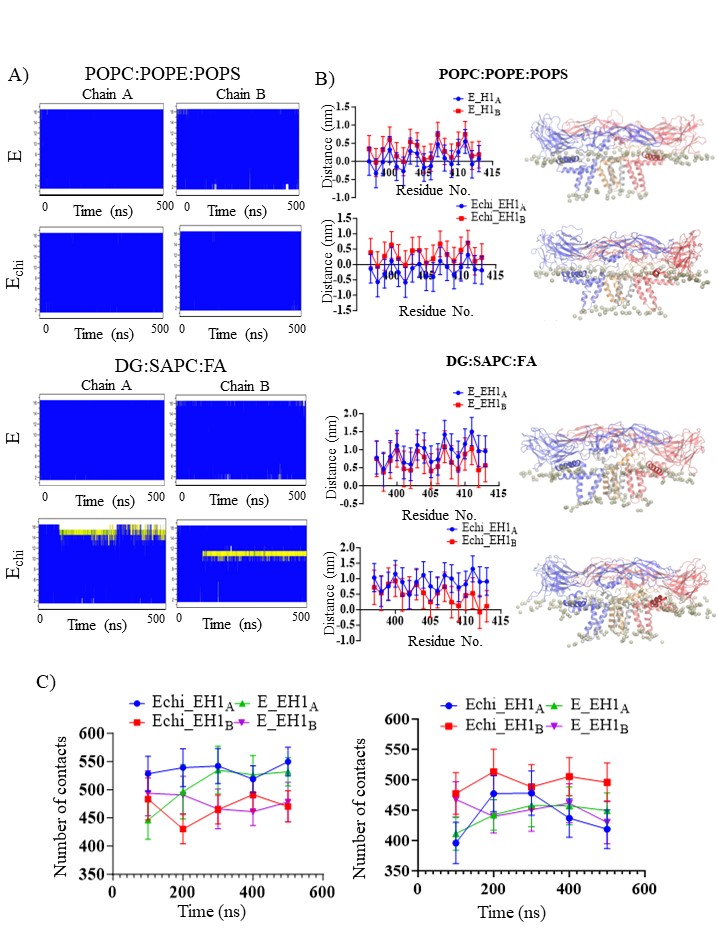

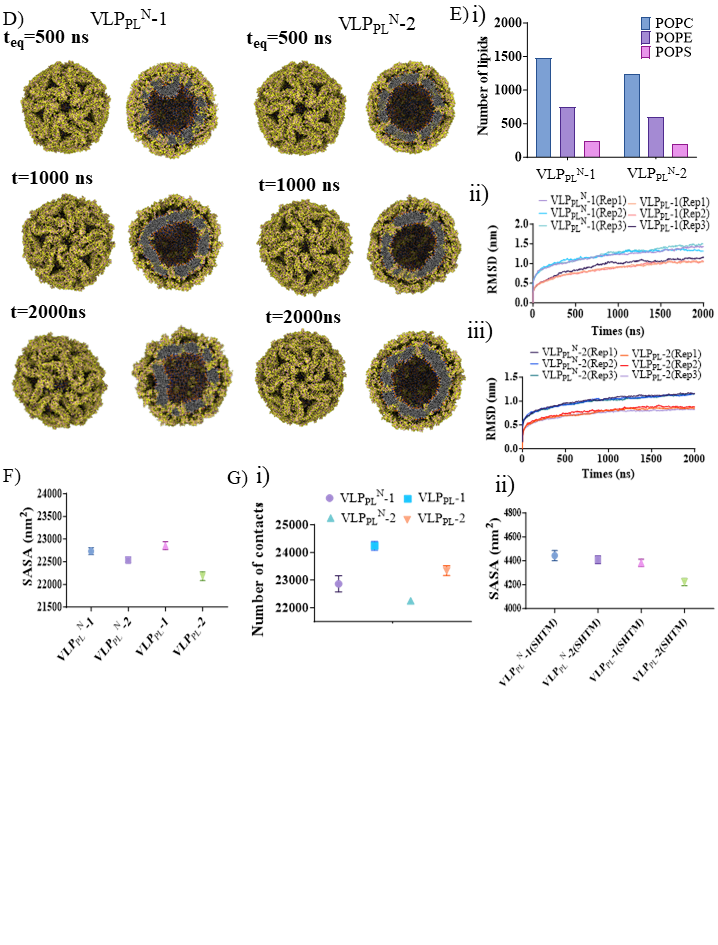

**Fig. S13. Comparing native DENV (VLP_PL_^N^) and chimeric DENV (VLP_PL_) VLP dynamics in PL dominant lipid envelope in CG simulations. (A)** Per-residue secondary structure changes over the simulation time for H1 helix (residues 397 to 415) from E and Echi dimer simulations. **(B)** Per-residue insertion of depth of E-H1 helix within lipid bilayers for PL dominant (top) and DG (bottom) dominant E/Echi-M protein simulations. Each point represents the average insertion depth with respect to lipid headgroups over final 200 ns of simulation. Alongside each plot, a snapshot of final coordinates of E-H1 is shown. E/M proteins are represented as transparent cartoon and lipid reference plane containing atoms are shown as transparent beads. E-H1 helices are shown in solid colors depicting the orientation with respect to membrane. Error bars were calculated as standard deviations over 500 ns of the trajectory. **(C)** EH1-helix-lipid contacts over the simulation time shown as 100 ns block averages for two lipid bilayers: PL (left) and DG (right). **(D)** Snapshots showing native DENV VLPs i.e., VLP_PL_^N -^1 and VLP_PL_^N^-2 CG structures from restrained equilibration (teq=500 ns) and unrestrained (t=1000 ns and t=2000 ns) simulation trajectories of length of 500 ns and 2000 ns respectively. **(E)** **(i)** Plot showing the number of lipid molecules in vesicles of native (VLP_PL_^N^-1 and VLP_PL_^N^-2) and chimeric (VLP_PL_-1 and VLP_PL_-2) DENV VLP CG models. **(ii-iii)** Comparison of RMSD of native and chimeric VLP CG simulations during 2000 ns long unrestrained production simulations. **(E)** Comparison of average solvent accessible surface (SASA) of protein shell averaged over the last 1000 ns of three replicate simulations from native (VLP_PL_^N^-1 and VLP_PL_^N^-2) and chimera (VLP_PL_-1 and VLP_PL_-2) VLP systems. **(F) (i)** Comparison of number of contacts between SH-TM region (residues 395-495) and lipid envelope from native (VLP_PL_^N^-1 and VLP_PL_^N^-2) and chimeric (VLP_PL_-1 and VLP_PL_-2) VLP simulations. Lipid molecules within a cutoff distance on 0.6 nm were considered as a contact with SH-TM and plots show average number of contacts from last 1000 ns of three replicate simulations. **(ii)** Plot showing mean SASA of SH-TM region from native (VLP_PL_-1 and VLP_PL_-2) and chimeric (VLP_PL_^N^-1 and VLP_PL_^N^-2) DENV VLPs. Mean and standard deviation of SASA correspond to last 1000 ns of simulation time from each of the three replicate simulations.

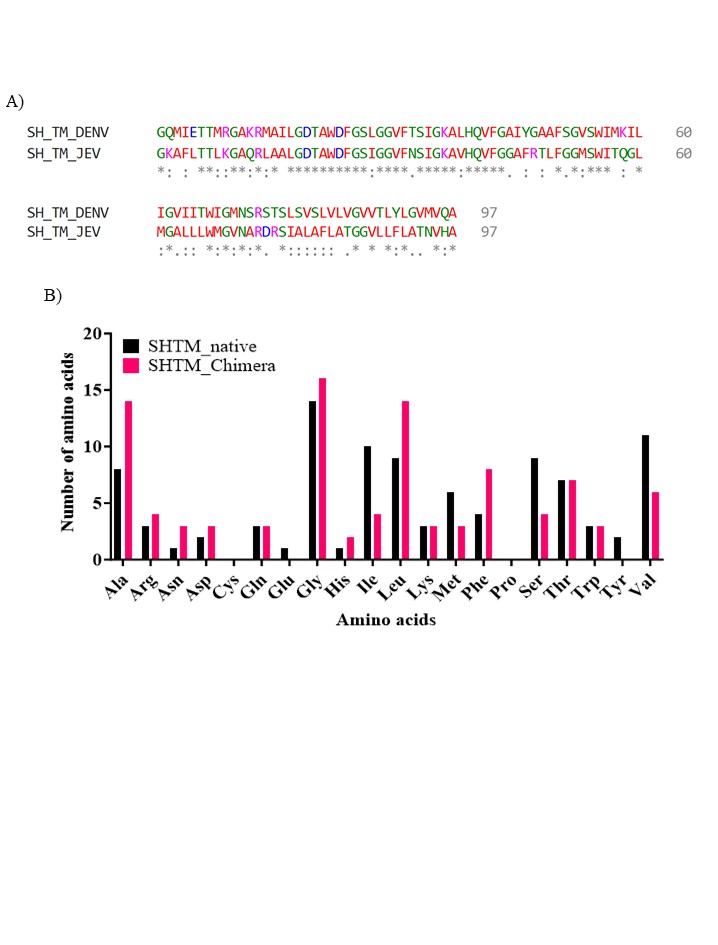

**Fig. S14.** **Amino acid composition of stem helix-transmembrane region (SHTM, residues 398-495). (A)** Sequence alignment of SHTM region from DENV and JEV E protein. (**B)** Plot comparing number of each of the 20 amino acids on SHTM region of DENV and JEV.

**Table S1. The specific amino acid SDM primer design for mD2VLP.**

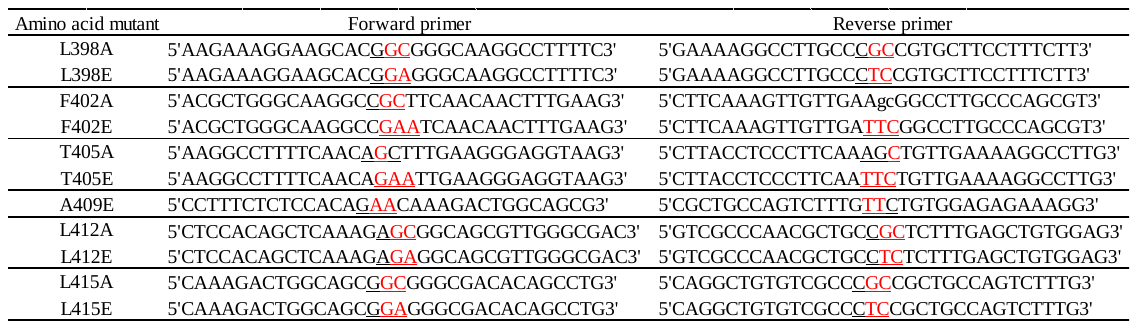

**Table S2. Summary of AA MD simulation system setups. Each E-M protein : lipid bilayer system was solvated in 66036 water molecules at 120mM NaCl concentration in a simulation box of 16.6×16.6×12.2 nm^3^ dimensions. Each system was subjected to 500 ns long MD simulation.**

|  | **E protein mutation** | **Lipid membrane**  **composition** |
| --- | --- | --- |
| DENV  (E_chi_-M protein) | WT | POPC:POPE:POPS  (60:30:10) |
|  | L398A |  |
|  | L398E |  |
|  | F402A |  |
|  | F402E |  |
|  | T405A |  |
|  | T405E |  |
|  | A409E |  |
|  | L412A |  |
|  | L412E |  |
| DENV  (E_chi_-M protein) | WT | DG:FA:SAPC  (56:26:8) |
|  | L398A |  |
|  | L398E |  |
|  | F402A |  |
|  | F402E |  |
|  | T405A |  |
|  | T405E |  |
|  | A409E |  |
|  | L412A |  |
|  | L412E |  |
| DENV | WT | POPC:POPE:POPS  (60:30:10) |

**Table S3.** **Summary of CG-MD simulation system setups.**

| **System** | **E protein mutation** | **Box Size (nm×nm×nm)** | **No. of water molecules** | **No. of lipid molecules** | **Simulation time (ns)** | **Lipid membrane**  **composition** |
| --- | --- | --- | --- | --- | --- | --- |
| **VLP_PL_** | VLP_PL_-1 | 40.5×40.5×40.5 | 507401 | 2486 | 2000 × 3 | POPC:POPE:POPS  (60:30:10) |
|  | VLP_PL_-2 | 32.5×32.5×32.5 | 237356 | 2055 | 2000 × 3 |  |
|  | VLP_PL_-3 | 32.5×32.5×32.5 | 239474 | 1737 | 2000 × 3 |  |
| **VLP_DG_** | VLP_DG_-1 | 32.5×32.5×32.5 | 235532 | 2886 | 2000 × 3 | DG:SAPC:FA  (56:26:8) |
|  | VLP_DG_-2 | 32.5×32.5×32.5 | 237188 | 2441 | 2000 × 3 |  |
|  | VLP_DG_-3 | 32.5×32.5×32.5 | 238864 | 2129 | 2000 × 3 |  |
| **VLP_PL_^N^** | VLP_PL_^N^-1 | 35.5×35.5×35.5 | 303266 | 2475 | 2000 × 3 | POPC:POPE:POPS  (60:30:10) |
|  | VLP_PL_^N^-2 | 34.5×34.5×34.5 | 276295 | 2039 | 2000 × 3 |  |

**Table S4**: Comparing the starting (t=0) and average values of physical properties of VLPs from CG-MD simulations

| **VLP system** | **Envelope dominant lipid** | **SASA (nm^2^)** | | | | | | | | | **Diameter across symmetric axis (nm at t=0)** | **Average diameter across symmetric axis (nm)** |
| --- | --- | --- | --- | --- | --- | --- | --- | --- | --- | --- | --- | --- |
|  |  | **D-I**  **(t=0)** | **D-I_AV_** | | **D-II**  **(t=0)** | **D-II_AV_** | | **D-III**  **(t=0)** | **D-III_AV_** | |  |  |
| VLP-1 | PL | 5789.8 | | 5864 | 7158.2 | | 7178 | 4571.7 | | 4669 | 26.4 | 25.6 |
|  | DG | 5742.9 | | 5813 | 7121.0 | | 7095 | 4549.3 | | 4568 | 26.3 | 23.7 |
| VLP-2 | PL | 5733.1 | | 5822 | 7103.9 | | 7132 | 4545.9 | | 4607 | 26.3 | 25.1 |
|  | DG | 5735.8 | | 5816 | 7113.4 | | 7087 | 4540.0 | | 4551 | 26.2 | 23.6 |
| VLP-3 | PL | 5728.1 | | 5801 | 7091.4 | | 7103 | 4527.8 | | 4581 | 26.2 | 24.0 |
|  | DG | 5709.6 | | 5807 | 7088.7 | | 7081 | 4539.6 | | 4561 | 26.2 | 23.5 |

**Movie S1.** 500 ns all atom simulation of Echi-M dimer embedded in phospholipid bilayer. Protein is shown in cartoon representation where E protein monomers are red and blue while M proteins are in grey and orange. Membrane phosphate groups are shown as brown spheres.

**Movie S2.** 500 ns all atom simulation of F402E mutant of Echi-M dimer embedded in phospholipid bilayer. Protein is shown in cartoon representation where E protein monomers are red and blue while M proteins are in grey and orange. Membrane phosphate groups are shown as brown spheres.

**Movie S3.** 500 ns all atom simulation of of F402L/A409L mutant of Echi-M protein dimer embedded in phospholipid bilayer. Protein is shown in cartoon representation where E protein monomers are red and blue while M proteins are in grey and orange. Membrane phosphate groups are shown as brown spheres.

**Movie S4.** 2000 ns long coarse-grained simulation trajectory of VLP_PL_ -1. VLP is shown in cross section where all beads are shown as spheres: protein backbone and side chain beads in orange and yellow respectively, lipids tails are shown in cyan while lipid head groups in blue, purple and brown.

**Movie S5.** 2000 ns long coarse-grained simulation trajectory of VLP_PL_ -2. VLP is shown in cross section where all beads are shown as spheres: protein backbone and side chain beads in orange and yellow respectively, lipids tails are shown in cyan while lipid head groups in blue, purple and brown.
